# *Starship* giant transposons dominate plastic genomic regions in a fungal plant pathogen and drive virulence evolution

**DOI:** 10.1101/2025.01.08.631984

**Authors:** Yukiyo Sato, Roos Bex, Grardy C. M. van den Berg, Monica Höfte, Michael F. Seidl, Bart P.H.J. Thomma

## Abstract

*Starships* form a recently discovered superfamily of giant transposons in Pezizomycotina fungi, implicated in mediating horizontal transfer of diverse cargo genes between fungal genomes. Their elusive nature has long obscured their significance, and their impact on genome evolution remains poorly understood. Here, we reveal a surprising abundance and diversity of *Starships* in the phytopathogenic fungus *Verticillium dahliae*. Remarkably, *Starships* dominate the plastic genomic compartments involved in host colonization, are enriched in virulence-associated genes, and exhibit genetic and epigenetic characteristics associated with adaptive genome evolution. We further uncover extensive horizontal transfer of *Starships* between *Verticillium* species and, strikingly, from distantly related *Fusarium* fungi. Finally, we demonstrate how *Starship* activity facilitated the *de novo* formation of a novel virulence gene. Our findings illuminate the profound influence of *Starship* dynamics on fungal genome evolution and the development of virulence.

## INTRODUCTION

Transposable elements (TEs, transposons) are ubiquitous mobile genetic elements in all life forms. Originally, these have been seen as selfish elements carrying only genetic information for their proliferation, but presently are appreciated to shape genome structure and function, and drive evolutionary innovations (Fedoroff, 2012). Whereas most TE superfamilies have simple structures and encode few proteins (Wicker et al., 2007), also “giant” TEs ranging from tens to hundreds of kilobases (kb) with tens to hundreds of “cargo” genes occur (Arkhipova and Yushenova, 2019; Inoue and Takeda, 2023; A. Urquhart et al., 2024).

*Starships* are giant TEs (15-700 kb) that were recently discovered in Pezizomycotina fungi, the largest subdivision of Ascomycota, and typically contain a tyrosine recombinase (YR) “captain” gene as first gene, required for transposition, while cargo genes are variable and functionally diverse (Gluck-Thaler et al., 2022; Gluck-Thaler and Vogan, 2024; A. Urquhart et al., 2024; Urquhart et al., 2023). How *Starships* impact global genome evolution, and the range and extent to which *Starships* have shaped genomes over time, remains enigmatic. *Starship* detection is technically challenging owing to the large diversity in cargo that can include abundant repeats (Gluck-Thaler et al., 2022), and relies on detection of presence/absence of YR-containing inserts in orthologous sites among highly contiguous genome assemblies (A. Urquhart et al., 2024). While 143 *Starships* were identified in this manner in a systematic search of 2,899 fungal genomes, 10,628 “orphan” YR genes remained, suggesting many overlooked *Starships* (Gluck-Thaler and Vogan, 2024). *Starships* occupy up to 2.4% of the genome of the human pathogen *Aspergillus fumigatus*, show extensive presence/absence variation, and contain many differentially expressed cargo genes upon infection (Gluck-Thaler et al., 2024). Moreover, in the plant pathogen *Macrophomina phaseolina*, 30% of chromosomal translocations, inversions, and putative chromosomal fusions occur near *Starship* insertions (Gluck-Thaler et al., 2022). Interestingly, *Starships* can transfer horizontally between closely-related fungi of the same order, and can transfer important traits such as pathogenicity (Bucknell et al., 2024; Bucknell and McDonald, 2023; Gourlie et al., 2022; O’Donnell et al., 2024; Peck et al., 2024; A. S. Urquhart et al., 2024; Urquhart et al., 2023, 2022).

Pathogens and their hosts typically engage in molecular arms races, with the pathogen exploiting secreted virulence factors (effectors) to mediate host colonization, while hosts employ immune receptors for pathogen interception (Cook et al., 2015; Dangl and Jones, 2024; Möller and Stukenbrock, 2017). To avoid recognition, pathogen effector catalogs are highly dynamic and variable, mediated by a “two-speed genome” organization in which virulence genes co-localize in highly plastic genomic regions that are enriched in repetitive elements and particular epigenetic features (Dong et al., 2015; Frantzeskakis et al., 2019; Kramer et al., 2023; Raffaele and Kamoun, 2012; Seidl et al., 2016; Seidl and Thomma, 2017; Torres et al., 2020). Accordingly, the fungus *Verticillium dahliae* that causes vascular wilt disease in hundreds of hosts (Fradin and Thomma, 2006; Inderbitzin et al., 2011; Klimes et al., 2015) contains plastic ‘adaptive genomic regions’ (AGRs) (Cook et al., 2020) that are enriched in virulence genes (Chavarro-Carrero et al., 2021; de Jonge et al., 2013, 2012; Kombrink et al., 2017; Snelders et al., 2023, 2020), transcriptionally active TEs (Cook et al., 2020; Faino et al., 2016; Torres et al., 2021), and structural variations (de Jonge et al., 2013; Faino et al., 2016; Klosterman et al., 2011; Torres et al., 2021), associated with a unique chromatin profile and physical interactions in the nucleus (Cook et al., 2020; Kramer et al., 2022, 2021; Torres et al., 2024). Besides *V. dahliae*, the Pezizomycotina *Verticillium* genus contains nine additional plant-associated species (Inderbitzin et al., 2011). Thus far, two *Starships* have been identified in a single strain of *V. dahliae* (Gluck-Thaler and Vogan, 2024). Here, we queried 56 highly contiguous *Verticillium* genome assemblies for *Starships* to analyze their association with the evolution of AGRs and virulence on plant hosts.

## RESULTS

### A wealth of *Starships* occurs in the *Verticillium* genus

To identify *Starships* across the *Verticillium* genus, we collected 56 high-quality *Verticillium* genome assemblies, comprising 36 *V. dahliae* strains and 20 strains of the nine remaining species (Fig. 1a; Supplementary Tables 1 and 2) and queried these genomes for *Starships* using “Starfish” (Gluck-Thaler and Vogan, 2024). We uncovered 54 *Starships* that belong to 24 haplotypes of 14 naves of 7 families (Fig. 1a-b; Supplementary Table 3). As between one and three *Starships* were detected in 33 of the 56 strains belonging to seven of the ten *Verticillium* species (Fig. 1a-b), most *Verticillium* genomes contain a *Starship*, and several genomes even multiple. Moreover, as these *Starships* range from 17 to 625 kilobases (kb) (Fig. 1c), and larger ones typically contain multiple YR genes of different naves (Fig. 1c-d), these likely represent nested *Starship* insertions. Thus, the final number of *Verticillium Starships* is likely under-estimated.

**Fig. 1.**
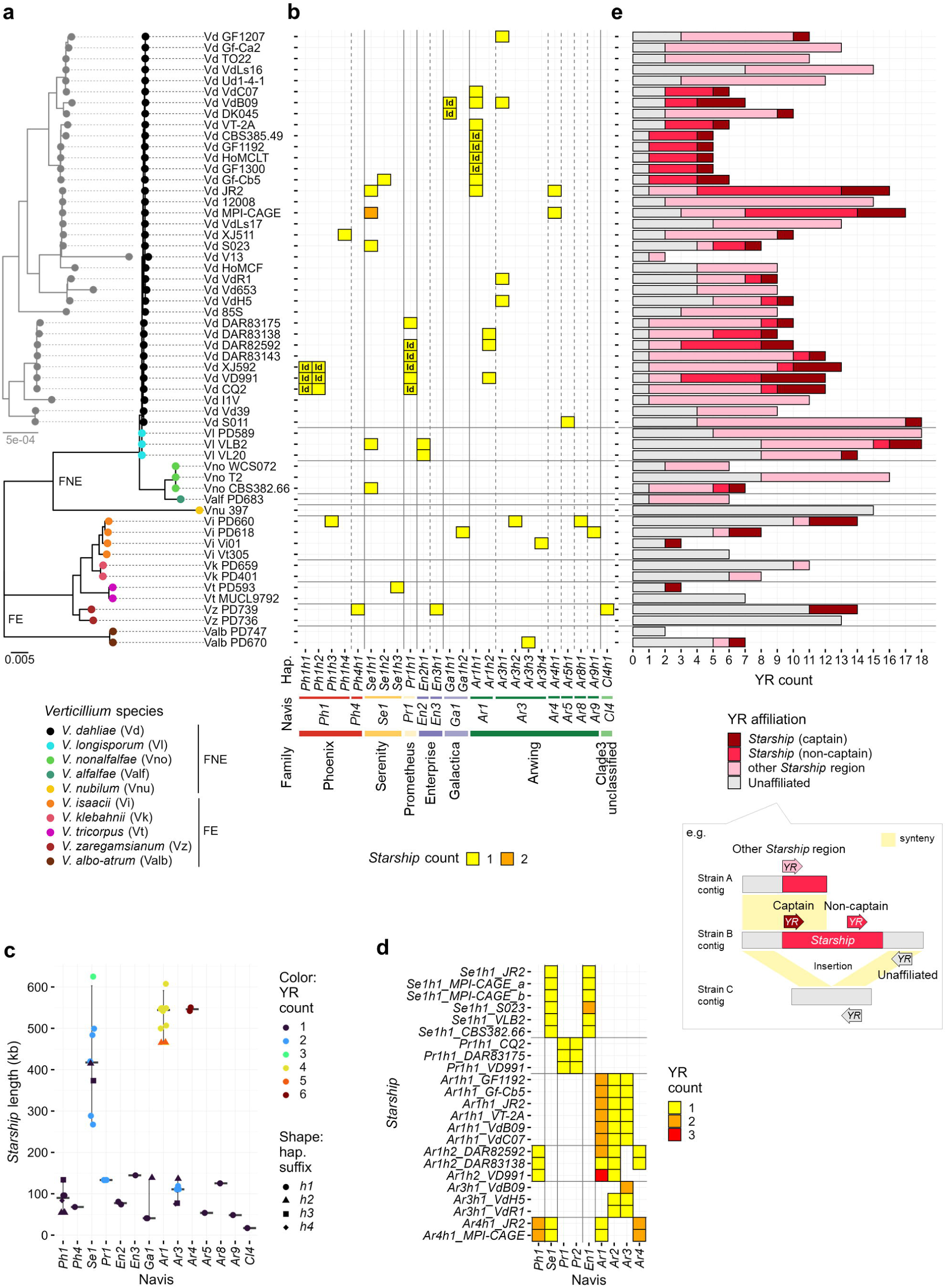
Diverse *Starships* populate the *Verticillium* genus. **a**, The tree in black shows the phylogeny of the 56 strains used in this study across the *Verticillium* genus, divided into the Flavexudans (FE) and Flavnonexudans (FNE) clades, based on whole-genome sequence alignments. Circle colors indicate the ten *Verticillium* spp. while the label comprises species abbreviation followed by strain name. The tree in grey shows only the *V. dahliae* strains at increased resolution. Scale bars indicate phylogenetic distances expressed as nucleotide substitutions per site. **b**, Repertoires of *Starship* haplotypes (hap.) per strain. *Starships* were classified according to previous studies (Gluck-Thaler and Vogan, 2024; A. Urquhart et al., 2024). Columns indicate *Starship* haplotypes defined by *k*-mer similarity and named according to captain navis and family (Extended Data Fig. 1b), whereas heatmap colors show haplotype member counts. “Id” refers to identical *Starships* (coverage and nucleotide sequence identity >98%) within a haplotype. **c**, Size of the different *Verticillium Starships*. Grey crossbars and error bars indicate the median and interquartile range of *Starship* lengths for each navis. **d**, YR navis repertoires in *Starships* with multiple YR genes. **e**, YR gene classification per strain where “*Starship* (captain)” indicates YR genes located as the first gene at the 5’-terminus of a *Starship* for which both borders could reliably be identified with the Starfish tool (Gluck-Thaler and Vogan, 2024), “*Starship* (non-captain)” refers to YR genes located at other sites in a *Starship*, suggesting nested *Starships* with unidentified boundaries. Furthermore, “other *Starship* region” indicates YR gene presence in regions that could not reliably be identified as *Starship* with Starfish, but that are syntenic to reference *Starships*. Finally, “unaffiliated” refers to YR genes that cannot reliably be affiliated with a *Starship* region.

*Verticillium* genomes exhibit extensive large-scale genomic rearrangements that may have affected the integrity of prior inserted *Starships* (de Jonge et al., 2013; Faino et al., 2016; Shi-Kunne et al., 2018). However, Starfish cannot identify fragmented *Starships*, rearranged *Starship* insertion sites, or *Starship* insertions into lineage-specific regions (Gluck-Thaler and Vogan, 2024). To identify such *Starships*, we queried for reference YRs previously identified in Pezizomycotina genomes (Gluck-Thaler and Vogan, 2024), revealing 2-18 homologs in each strain, amounting to 556 YR genes of 38 naves and seven families (Extended Data Fig. 1 and Supplementary Table 4). Only 21% of them belong to *Starships* identified by Starfish (Fig. 1e), while the remaining 79% could point to unidentified *Starships*. Additionally, we identified syntenic regions to *Starships* as “*Starship* regions”. Intriguingly, such *Starship* regions occur in all strains (Supplementary Table 5) and half of the YR genes that did not occur in *Starships* appeared in such *Starship* regions (Fig. 1e). Thus, we reveal abundant *Starships* and their remnants in the *Verticillium* genus.

### *Starships* are hotspots of large-scale genomic rearrangements

To detail how genomic rearrangements affected *Starships,* we compared telomere-to-telomere genome assemblies of *V. dahliae* strains JR2 and VdLs17 that comprise dozens of large-scale genomic rearrangements (de Jonge et al., 2013; Faino et al., 2016, 2015). In strain JR2, we detected three large *Starships* of 0.50-0.54 Mb each, belonging to haplotypes *Ar1h1*, *Ar4h1*, and *Se1h1*, plus additional *Starship* regions, collectively accounting for 5.3% (1.92 Mb) of the genome (36.15 Mb) (Fig. 2a). Although no *Starships* were detected in VdLs17 by Starfish, *Starship* regions account for 2.7% (0.96 Mb) of the genome (35.97 Mb) (Fig. 2a). Intriguingly, 60% of the inter-chromosomal rearrangement breakpoints between these strains occurred in *Starship* regions (Fig. 2a). Accordingly, *Starship* regions were mainly detected in AGRs that are enriched in such rearrangements(de Jonge et al., 2013; Faino et al., 2016) (Fig. 2a-b). In strain JR2, 92% (1.77 Mb) of the *Starship* regions colocalized with AGRs (Supplementary Table 6). Moreover, 53% of the total AGR complement belongs to *Starships*. In VdLs17, 94% (0.90 Mb) of the *Starship* regions colocalized with 20% of the AGR complement (Supplementary Table 6).

**Fig. 2.**
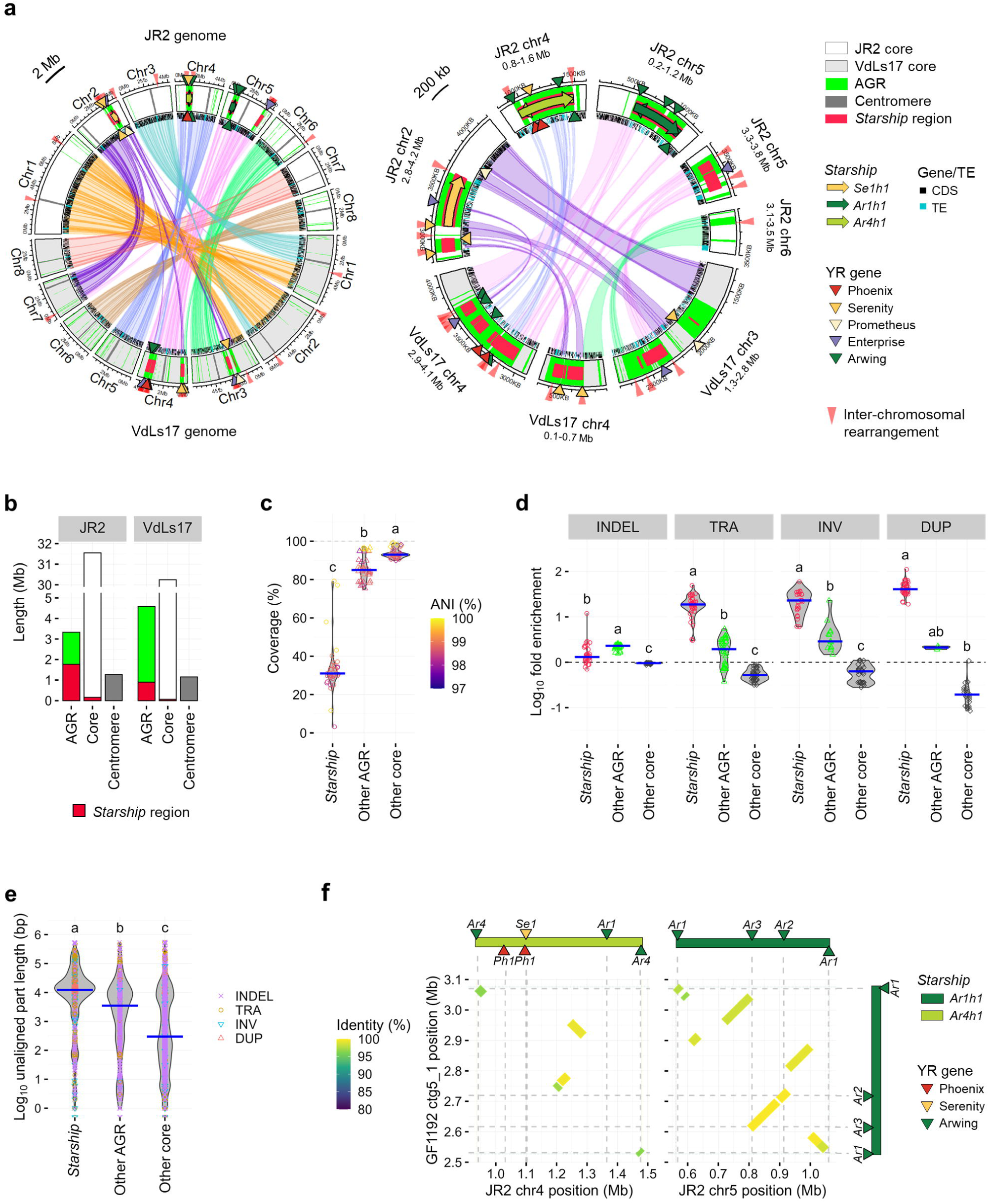
*Starships* are hotspots of genomic rearrangements. **a**, Circular plots showing the locations of *Starship* regions, AGRs, and genomic rearrangements between *V. dahliae* strains JR2 (upper eight chromosomes) and VdLs17 (lower eight chromosomes). Tracks are filled with colors representing either core, AGR, or centromeric regions, overlaid with bold lines and arrows representing *Starship* regions and *Starships* colored by haplotype. Regular triangles point to YR genes colored by family and annotated with navis ID. Colored bands at the inner edge of tracks represent genetic elements grouped into protein coding sequence (CDS) and transposable element (TE). Ribbons connect syntenic regions (>80% nucleotide sequence identity over 10 kb). **b**, Total length of genomic compartments in *V. dahliae* strains JR2 and VdLs17. **c**, Violin plots depicting the sequence alignment coverage determined by comparing JR2 genomic compartments against 35 *V. dahliae* genomes (Extended Data Fig. 2). Points indicate the coverage and the color represents genome-wide Average Nucleotide Identity (ANI), while blue crossbars indicate the median values. Different letter labels indicate significant differences (Dunn’s test, *p* < 0.05). **d**, Violin plots depicting the fold-enrichment of structural variations (SVs) (insertion and deletion (INDEL) without size definition, translocation (TRA), inversion (INV), and duplication (DUP)) in the three genomic compartments compared with the genome-wide average. SVs were determined by comparing the genome of *V. dahliae* strain JR2 against the 35 *V. dahliae* genomes. Points indicate the fold-enrichment determined for each genome, while blue crossbars indicate the median values. Different letter labels indicate significant differences for each genetic variation. **e**, Violin plots depicting length of unaligned regions accompanied by each SV in JR2 genomic compartments. Points indicate the length of every gap found between JR2 genome and 35 *V. dahliae* genomes in a color and shape representing SV type, while blue crossbars indicate the median values. Different letter labels indicate significant differences (Dunn’s test, *p* < 0.05). **f**, *Starship* rearrangements between the genomes of *V. dahliae* strains JR2 (X-axis) and GF1192 (Y-axis). Diagonal lines indicate synteny while the color represents nucleotide sequence identity. Bars and triangles aligned to the plots indicate the positions of *Starships* and YR genes.

Since significant AGR proportions could not be assigned to *Starships* in JR2 (47%) and VdLs17 (80%), we determined whether *Starship* regions in AGRs display different characteristics than other AGRs. Pairwise genomic alignments of the JR2 genome with those of the other 35 *V. dahliae* strains revealed significantly lower alignment coverage in *Starship* regions (median 31%) than in other AGRs (median 85.5%) and core genomic regions (median 93%) (Fig. 2c and Extended Data Fig. 2), indicating enhanced presence/absence variation in *Starship* regions. Moreover, we revealed an extreme enrichment in structural variation, particularly concerning translocations, inversions, and duplications, in *Starships* (Fig. 2d and Supplementary Table 7). Regions lacking alignment accompanying these rearrangements were significantly larger in *Starship* regions (median 12.2 kb) than in other AGRs (median 3.4 kb) and core regions (median 0.3 kb) (Fig. 2e), underscoring the extensive presence/absence variation in *Starship* regions.

Intriguingly, some genomic rearrangements even occurred between *Starships* as the *Ar1h1 Starship* in *V. dahliae* strain GF1192 is largely syntenic to the JR2 *Ar1h1 Starship*, but synteny lacks at four sites that are syntenic to parts of the JR2 *Ar4h1 Starship* (Fig. 2f), leading to *Starship* diversification.

### *Starships* display typical traits of plastic genomic regions

Like plastic genomic regions of many filamentous plant pathogens (Dong et al., 2015; Kramer et al., 2023; Raffaele and Kamoun, 2012; Seidl et al., 2016; Seidl and Thomma, 2017), *V. dahliae* AGRs are enriched in effector genes, *in planta* induced genes with facultative heterochromatic histone modifications (H3K27me3) (Cook et al., 2020; de Jonge et al., 2013; Kramer et al., 2022), and repetitive elements including active TEs (Faino et al., 2016; Torres et al., 2021). Intriguingly, the previously characterized effector genes *Ave1* (de Jonge et al., 2012; Snelders et al., 2020) and *Av2* (Chavarro-Carrero et al., 2021) occur in *Ar1h1* and *Ar4h1 Starships* (Fig. 3a-b). While no difference in *in planta* gene induction and H3K27me3 levels could be observed between *Starship* regions and other AGRs (Fig. 3c-d and Supplementary Table 8), the average TE density and expression were higher in *Starship* regions than in other AGRs (Fig. 3e-f and Supplementary Table 9). *V. dahliae* AGRs are furthermore characterized by segmental duplications that physically interact in the nucleus (Faino et al., 2016; Torres et al., 2024). Interestingly, consistent with the segmental duplication pattern (Fig. 2d and Fig. 3g), such bipartite long-range chromatin interactions occur between the *Ar1h1* and *Ar4h1 Starships*, between nested *Starships* within the *Ar4h1 Starship*, and between the *Se1h1 Starship* and syntenic *Starship* regions in the JR2 genome (Fig. 3h). Collectively, our data indicate that *Starships* in *V. dahliae* are strongly associated with traits that characterize AGRs.

**Fig. 3.**
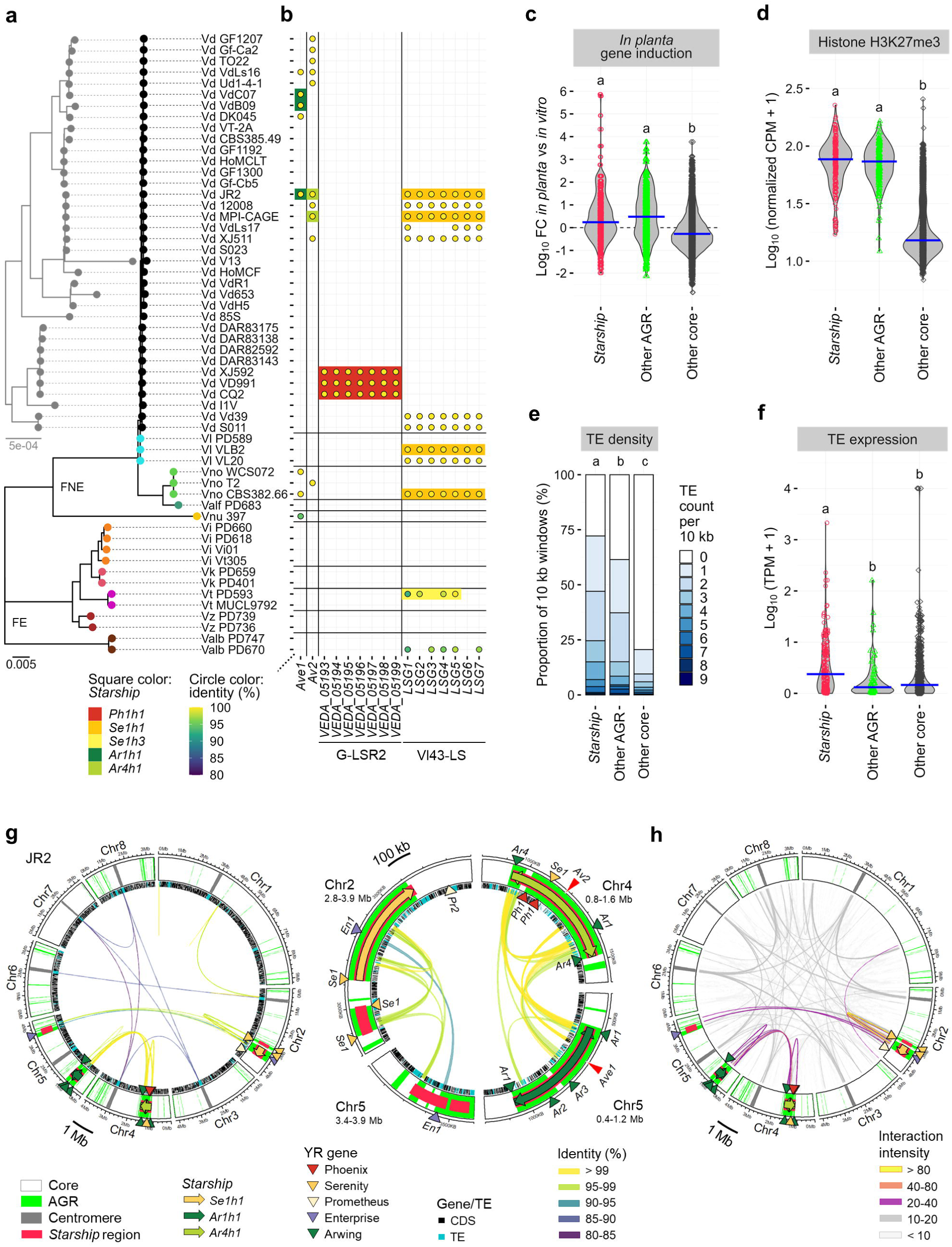
*Starships* show typical traits of the plastic genomic regions. **a**, *Verticillium* phylogeny as described in Fig. 1a. **b**, Presence or absence of (a)virulence-associated genes in the 56 *Verticillium* strains. Square colors indicate *Starships* in which the genes were detected. Circles indicate the presence of orthologs with 100% gene coverage and the fill color representing nucleotide identity. **c**, Violin plots depicting the *in planta* expression induction of genes residing in the three genomic compartments in *V. dahliae* strain JR2. Induction levels are represented by fold-changes (FC) of gene expression *in planta* (*Arabidopsis thaliana*) versus *in vitro* (potato dextrose broth, PDB). Points indicate FC values for each gene, while blue crossbars indicate median values. Different letter labels indicate significant differences (Dunn’s test, *p* < 0.05). **d**, Violin plots depicting histone H3K27me3 levels for the three JR2 genomic compartments expressed as ChIP-Seq read counts per million (CPM) normalized by bin length. Each point indicates the value for a bin (∼10 kb), while blue crossbars indicate median values. Different letter labels indicate significant differences (Dunn’s test, *p* < 0.05). **e**, Stacked bar plots depicting the transposable element (TE) density in the three JR2 genomic compartments over 10 kb windows. **f**, Violin plots depicting the TE expression levels in the three JR2 genomic compartments expressed in transcripts per million (TPM) for *V. dahliae* cultivated in PDB. Points indicate TPM values for individual TEs, while blue crossbars indicate median values. Different letter labels indicate significant differences (Dunn’s test, *p* < 0.05). **g**, Locations of *Starships* and AGRs in the JR2 genome. See Fig. 2a legend for the details of symbols. Ribbons connect pairs of segmentally duplicated regions that share >80% nucleotide sequence identity over 10 kb, with a color representing identity. **h**, Long-range chromatin interactions in the JR2 genome. Ribbons connect genomic regions that are separated in the genome but physically interact in nuclei by colors that represent the intensity.

### *Starships* carry virulence cargo genes

To assess whether besides *Ave1* and *Av2* that occur in *Ar1h1* and *Ar4h1 Starships*, further previously characterized virulence genes appear as cargo, we queried the *Starships* for *Verticillium* genes that are described in the Pathogen-Host Interactions Database (PHI-base) (Urban et al., 2022). This revealed that the G-LSR2 region, proposed to confer cotton-specific virulence in *V. dahliae* (J.-Y. Chen et al., 2018), occurs in a *Ph1h1 Starship*, while the Vl43-LS region that affects *V. longisporum* virulence (Harting et al., 2021) occurs in a *Se1h1 Starship* (Fig. 3a-b). In addition, several *Starships* (*Se1h1*, *Se1h2*, *Se1h3*, and *En3h1*) contain orthologs of other virulence-associated genes, encoding transporters, β-tubulin, and regulators of infection structure morphogenesis and pH-signaling (Caracuel et al., 2003; Kodama et al., 2017; Nguyen et al., 2008; Ortoneda et al., 2004; Zhang et al., 2017) (Extended Data Fig. 3 and Supplementary Table 10), demonstrating that *Verticillium Starships* carry diverse virulence-associated cargo genes.

### *Starship* dynamics within and between *Verticillium* genomes

HGT among *Verticillium* spp. may have contributed to the shaping of AGRs (Depotter et al., 2019). To detect possible *Starship* transfer between *Verticillium* spp., we utilized an implicit phylogenetic method that identifies segments with increased sequence similarity over the genome-wide average nucleotide identity (ANI) (Ravenhall et al., 2015). This analysis revealed that several *Starships*, such as those belonging to haplotypes *Se1h1* and *Ar1h1*, as well as syntenic *Starship* regions have higher ANI levels than the genome-wide average between the *Verticillium* species in which they occur (Fig. 4a-b, Supplementary Tables 11 and 12). The most conspicuous is the 0.5 Mb *Ar1h1 Starship* that carries *Ave1* and shares over 99% identity between *V. dahliae* and *V. nonalfalfae,* while the genome-wide average is only 93% (Fig. 4c, Supplementary Tables 11 and 12).

**Fig. 4.**
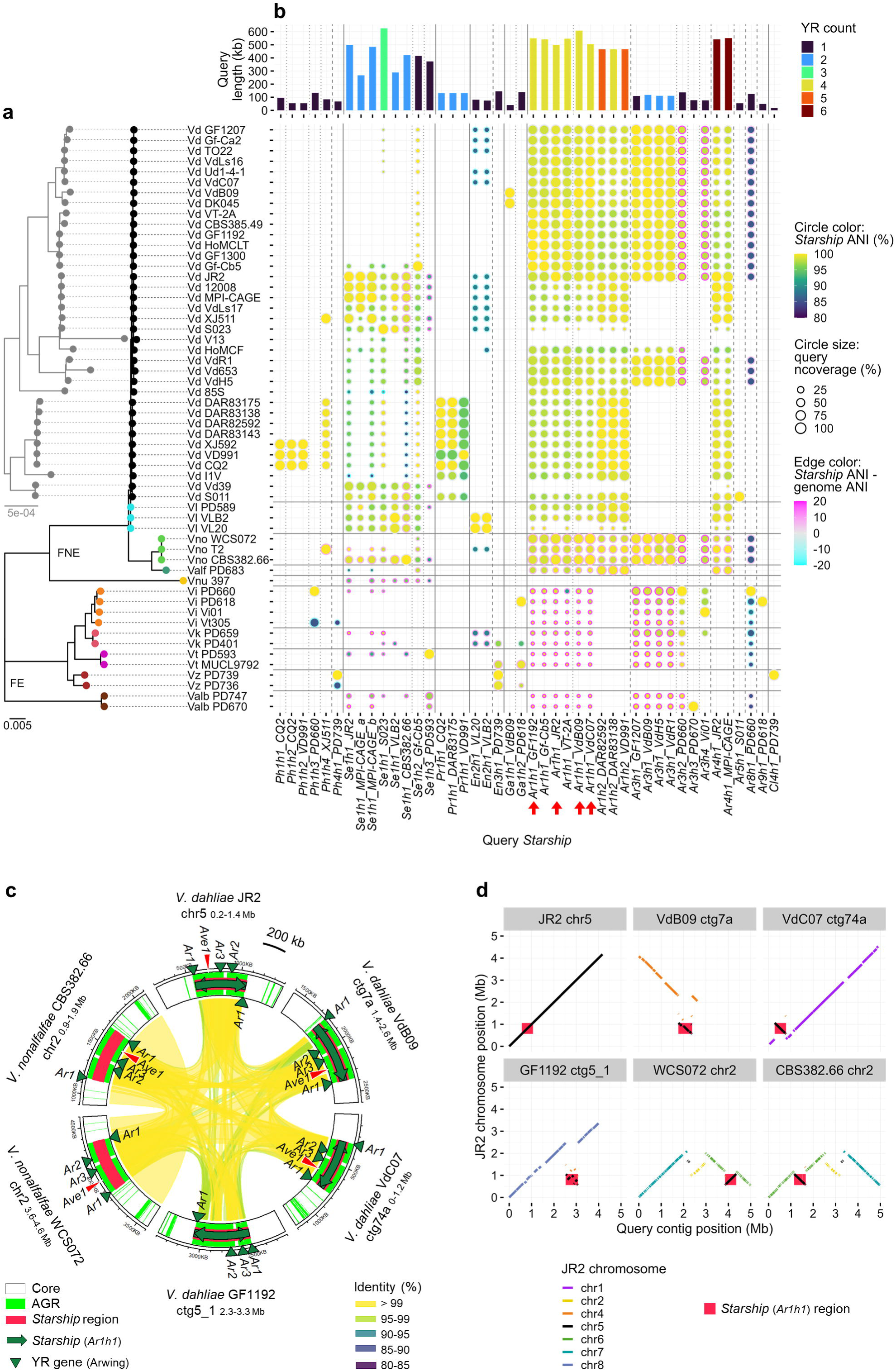
*Starship* mobility among *Verticillium* genomes. **a**, *Verticillium* phylogeny as described in Fig. 1a. **b**, Occurrence of diverse *Starships* across the *Verticillium* genus. Columns indicate different *Starships* with their length and number of YR genes. The heat map indicates the query coverage, ANI of *Starship* regions (*Starship* ANI), and the difference between *Starship* ANI and genome-wide ANI for each pairwise strain comparison to detect potential horizontal transfers. Red arrows indicate the *Starship* highlighted in (**c**). **c**, Circular plot depicting the synteny among regions of four *V. dahliae* and two *V. nonalfalfae* strains harboring orthologous *Ar1h1 Starships*. See Fig. 2a legend for the details of symbols. **d**, Synteny plots between the *V. dahliae* JR2 genome and the *Ar1h1 Starhip* regions in (**c**). Each plot indicates the synteny between the JR2 chromosomes (Y-axis) and each chromosome (chr) or contig (ctg) containing the *Ar1h1 Starship* (X-axis), with diagonal lines colored by syntenic JR2 chromosome. Red background indicates the coordinates of *Ar1h1 Starship*.

To further support *Starship* mobility, we compared synteny of orthologous *Starship* flanking regions, revealing that *Se1h1* and *Ar1h1 Starships* occur in several non-homologous genomic regions in different *Verticillium* clades (Fig. 4c-d and Extended Data Fig. 4a-b). For instance, the *Ar1h1 Starship* that occurs in nine *V. dahliae* strains and in two *V. nonalfalfae* strains collectively inserted into regions that are orthologous to five different JR2 chromosomes (Fig. 4d), suggesting at least five independent *Ar1h1 Starship* insertions.

Comparisons among orthologous *Starship* regions across *Verticillium* strains revealed various diversification patterns (Extended Data Fig. 4a-b). Besides nested *Starship* insertions (e.g. *Ar3h1 Starship* in *Ar1h1 Starship*, Extended Data Fig. 4c; Fig. 1c-d), we typically observed gain and loss of cargo elements in TE-enriched *Starships* (e.g. *Pr1h1 Starships*, Extended Data Fig. 4d). We furthermore observed invasion of orthologous sites by different *Starships* that contain orthologous captains, but otherwise lack synteny (e.g. *Ar1h1* and *Ar1h2 Starships*, Extended Data Fig. 4e), which points towards independent insertions as orthologous captains target particular sequences as *Starship* insertion sites (Gluck-Thaler and Vogan, 2024; Urquhart et al., 2023; Vogan et al., 2021).

### Cross-order horizontal *Starship* transfer

To detect possible horizontal *Starship* transfer between *Verticillium* and other fungi, we queried all Pezizomycotina genomes available in the GenBank WGS database (10,113 genomes from 668 genera, Extended Data Fig. 5a and Supplementary Table 13) with *Verticillium Starships*. Intriguingly, regions syntenic to *Ph1h1* and *Ph1h2 Starships* that were detected in only three *V. dahliae* strains (CQ2, VD991, and XJ592) (Fig. 4a-b), were found in the genomes of various *Fusarium* species, some of which contain regions syntenic to the G-LSR2 region (*Ph1h1 Starship* cargo) that was associated with *V. dahliae* virulence (J.-Y. Chen et al., 2018) (Fig. 5a and Extended Data Fig. 5b). Since *Fusarium* belongs to Hypocreales and synteny to these *Verticillium Starships* lacks in non-*Verticillium* genomes of the Glomerellales to which *Verticillium* spp. belong, these results suggest horizontal transfer of *Ph1h1* and *Ph1h2 Starships* between *Verticillium* and *Fusarium*.

**Fig. 5.**
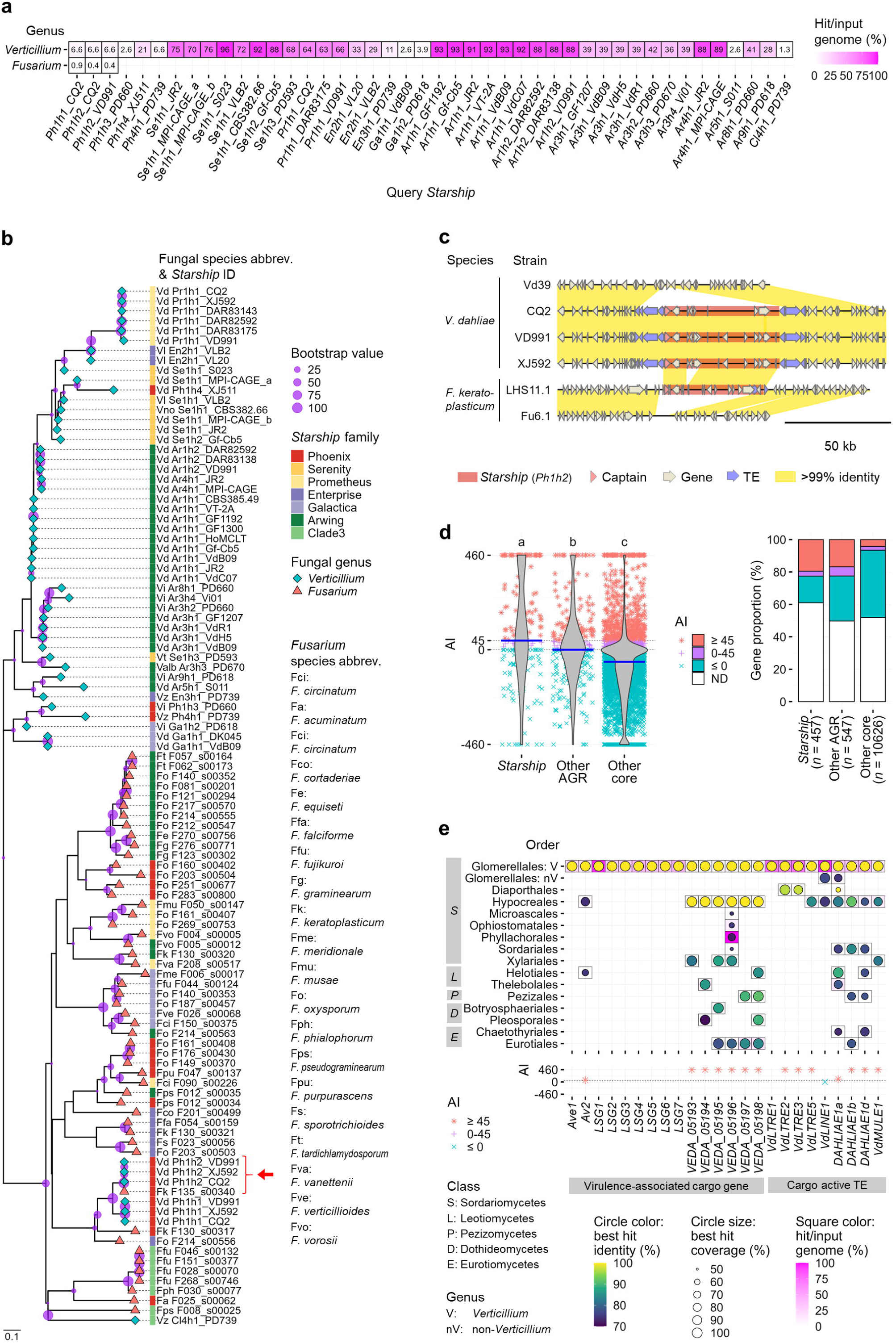
*Starship* dynamics among fungal orders. **a**, Occurrence of *Verticillium Starships* across 10,113 Pezizomycotina genomes. Cell colors and labels denote the percentage of genomes with hits (e-value <0.05 and total query coverage >50% or >30 kb) in each genus. **b**, Phylogeny of *Starships* that occur in the *Verticillium* and *Fusarium* genera based on *k*-mer similarity. Scale bar indicates the Mash distance that represents *k*-mer difference (Ondov et al., 2016). Circles at the nodes represent bootstrap values for 1,000 iterations. Red arrows indicate the *Starship* highlighted in (**c**). **c**, Similarity of *Ph1h2 Starships* and surrounding regions between *V. dahliae* and *F. keratoplasticum.* **d**, Possibility of horizontal gene transfer (HGT) between *Verticillium* and fungi belonging to the other orders of Pezizomycotina for genes residing the three genomic compartments in *V. dahliae* strain JR2. High Alien index (AI) values (≥45) suggest HGT, while moderate values (>0 and <45) suggest a weak HGT possibility. Points in the left violin plots indicate AI values of individual genes, while blue crossbars indicate median values. The right bar plots depict the percentage of genes with high, moderate, and low AI values. ND indicates that AI was not determined because of lacking hits in both ingroup and outgroup. **e**, Occurrence of homologs of *Verticillium Starship* cargo genes and TEs across 10,113 Pezizomycotina genomes. Square colors represent the percentage of genomes with hits (e-value <0.05, over 70% nucleotide identity over 50% coverage) in each group. Circles indicate the coverage and identity of the best hits. The lower panel indicates AI values as described in (**d**).

To identify the directionality of horizontal transfer, we analyzed a *k*-mer-based phylogeny of *Starships* (Hill et al., 2024b). To this end, we identified 52 *Starships* in 283 high-quality *Fusarium* genome assemblies (Supplementary Tables 14 and 15) and analyzed their phylogeny together with the 54 *Verticillium Starships*, revealing that most *Starships* clustered according to fungal phylogeny (Fig. 5b). In contrast, the *Verticillium Ph1h1* and *Ph1h2 Starships* phylogenetically localized in the *Fusarium Starship* clade (Fig. 5b), suggesting *Fusarium* to *Verticillium* transfer. Moreover, we identified a near-identical *Ph1h2 Starship* that shares over 99% nucleotide identity over 50 kb in three *V. dahliae* strains and in *F. keratoplasticum* (Fig. 5c). The *Starship* flanking regions are syntenic in three phylogenetically clustered *V. dahliae* strains, suggesting that the *Ph1h2 Starship* invaded *V. dahliae* from *F. keratoplasticum* before these strains diverged (Fig. 5c).

To further explore potential *Starship*-associated HGTs overlooked by the synteny search, we tested HGT signatures for each *Starship* cargo element using the Alien index (AI) that detects higher similarity to outgroup than ingroup orthologs (Gladyshev et al., 2008). We queried orthologs of *Verticillium* genes in the Pezizomycotina genomes and calculated AI scores with non-*Verticillium* Glomerellales genera as ingroup and other 74 orders as outgroup. This showed that the median AI for JR2 genes is higher in *Starship* regions (45) than in other AGR (0) and core regions (-59), and that the proportion of genes with AI ≥ 45 (indicative of HGT) is higher in *Starship* regions (19%) and other AGRs (17%) than in core regions (4%), suggesting cross-order HGT in *Starship* regions and other AGRs (Fig. 5d and Supplementary Table 16). Among the virulence-associated cargo genes, *Av2* as well as the G-LSR2 genes showed AI > 45 with hits in the various *Fusarium* spp. of the Hypocreales while lacking ingroup hits (Fig. 5e and Supplementary Table 17). Horizontal *Av2* transfer was further supported by phylogenetic analyses in which *Verticillium* orthologs were detected in a clade within a larger clade of *Fusarium* orthologs (Extended Data Fig. 6a-b). *Fusarium Av2* orthologs were found proximal to YR genes, which suggests *Starship* association (Extended Data Fig. 6c). We also tested the HGT signature of transcriptionally active *Verticillium* TEs (Amyotte et al., 2012) that occur in *Starship* regions (Extended Data Fig. 7a). Surprisingly, seven out of nine cargo TEs showed AI > 45 with the best outgroup hits in five orders including Hypocreales, while six of them lacked ingroup hits (Fig. 5e and Supplementary Table 17). Orthologs of the five cargo TEs also occurred in *Fusarium Starships* (Extended Data Fig. 7b), suggesting horizontal transfer via *Starships*. Collectively, our results provide evidence for horizontal transfer of various *Starship* cargo elements between *Verticillium* and fungi that even belong to other orders.

### *Starships* mediate *de novo* gene birth

*Starships* carry diverse lineage-specific genes (Gluck-Thaler et al., 2022), yet their origin remains enigmatic. We pursued the evolution of the *NLP6* effector gene that occurs in a limited number of *V. dahliae* strains, including VdLs17 (Santhanam et al., 2013), and was detected near a *Starship* region (Extended Data Fig. 8a). NLP6 belongs to the family of necrosis-and ethylene-inducing peptide 1 (Nep1)-like proteins (NLPs), many members of which confer virulence (Santhanam et al., 2013; Seidl and Ackerveken, 2019). As multiple NLP paralogs occur in *Verticillium* genomes (Santhanam et al., 2013), we first analyzed the phylogeny of NLP homologs in the 56 *Verticillium* genomes. NLP6 occurs in the same clade as NLP3 orthologs which were detected in all 56 strains (Fig. 6a-c). Intriguingly, NLP6 is more closely related to NLP3 orthologs of *V. nubilum* and species of the FE clade than to those of *V. dahliae* or other species of the FNE clade (Fig. 6a). The similarity between NLP3 orthologs and NLP6 occurs at the C-but not at the N-terminus (Extended Data Fig. 8b). Rather, the 3’ end (115-450 nt) of *NLP6* shares 90% nucleotide identity with the 3’ end (379-714 nt) of *V. nubilum NLP3*, which greatly exceeds the genome average (82%) (Fig. 6d, Extended Data Fig. 8c, and Supplementary Tables 12 and 18), suggesting that the 3’ part of *V. dahliae NLP6* is derived from a horizontally transferred *NLP3* ortholog. To explore the origin of the 5’ part (1-114 nt) of *NLP6*, we queried the *Verticillium* genomes and detected hits with 96% nucleotide identity in non-coding regions in *Se1h1 Starships* in four *Verticillium* species, which we termed *tNLP6* (for *truncated NLP6*) (Fig. 6d-f and Supplementary Table 18). Intriguingly, three *tNLP6* copies were detected around *NLP6* in VdLs17 (Fig. 6e). Collectively, these results suggest that *NLP6* was formed through the fusion of a non-coding *Starship* element with the 3’ part of an *NLP3* ortholog through a genomic rearrangement (Fig. 6g). Interestingly, although a role of *NLP3* in fungal virulence could not be demonstrated (Santhanam et al., 2013), deletion of *NLP6* results in reduced symptom development in *V. dahliae*-inoculated tomato plants (Fig. 6h and Extended Data Fig. 8d-e). Collectively, these results demonstrate that *Starships* mediated the emergence of a virulence-associated lineage-specific effector gene in *V. dahliae*.

**Fig. 6.**
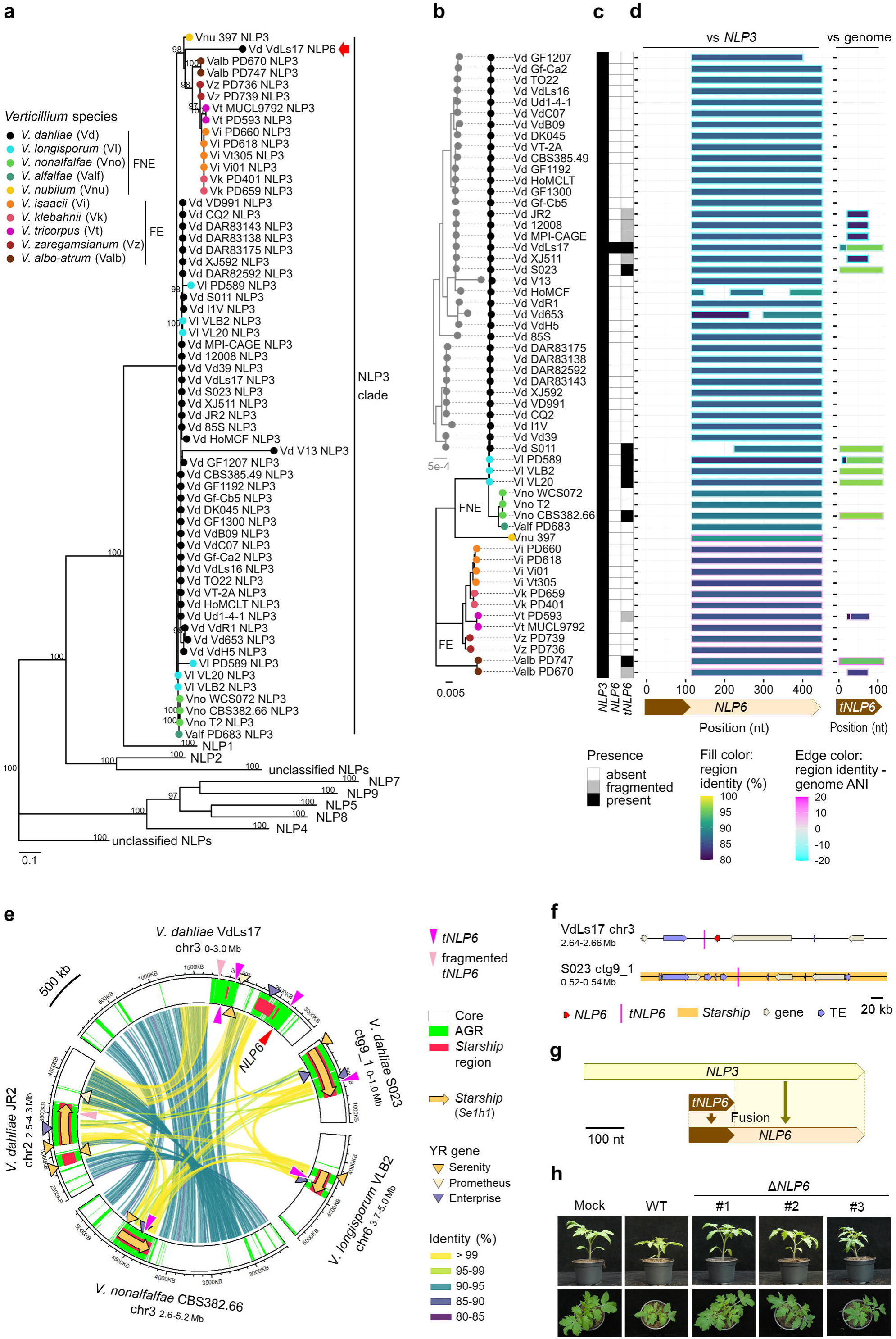
*Starships* contributed to *de novo* formation of the virulence gene *NLP6*. **a**, Phylogeny of necrosis and ethylene-inducing peptide 1 (Nep1)-like proteins (NLPs) in 56 strains across the *Verticillium* genus with all NLP clades collapsed except for the NLP3 clade. The red arrow points to NLP6. Scale bar indicates amino acid substitutions per site. Bootstrap values (>95%) for 1,000 iterations are shown at the nodes. **b**, *Verticillium* phylogeny as described in Fig. 1a. **c**, Presence or absence of *NLP3*, *NLP6*, and a truncated *NLP6* (*tNLP6*: 1-114 nt of *NLP6*) in the *Verticillium* strains. **d**, Coverage plots for alignments of *NLP6* with *NLP3* genes (left) and for alignments of *tNLP6* with genome sequences (right). The X-axes represent the nucleotide positions in *NLP6* or *tNLP6* of *V. dahliae* VdLs17. Bar colors indicate the identity of the region and its difference from genome-wide ANI between each *Verticillium* strain and VdLs17. **e**, Location of *tNLP6* in *Starship* in three *Verticillium* species. See Fig. 2a legends for the details of symbols. **f**, Location of *tNLP6* in non-coding regions of *V. dahliae* strains. **g**, Proposed model for *NLP6* evolution. **h**, Symptoms of tomato plants inoculated with wild-type (WT) and three independent *NLP6* deletion (Δ) lines of *V. dahliae* strain VdLs17 at 14 days post inoculation.

## DISCUSSION

Here, we reveal the unprecedented impact of *Starships* on fungal genome evolution through four main findings. First, every *Verticillium* genome contains *Starships* or their remnants. Secondly, these *Starships* compose the vast majority of AGRs that govern pathogenicity on plant hosts. Thirdly, extensive horizontal *Starship* transfer occurred between *V. dahliae* and phylogenetically diverse fungi. Fourthly, *Starships* contributed to the *de novo* generation of a novel virulence gene. Thus, *Starships* are not only instrumental for *Verticillium* genome evolution, but also fundamental to evolutionary innovations that led to plant pathogenicity.

Thus far, little is known about *Starship* abundance in individual fungal genomes. We identified 24 *Starship* haplotypes in 56 *Verticillium* genomes. These *Starships* generally occur in multiple *Verticillium* strains, indicating invasion before strain diversification. While some remained intact, others got disrupted by genomic rearrangements. The abundance in *Starships* exceeds the abundance previously recorded in *A. fumigatus*, where 20 *Starships* and their remnants were identified in a survey of 519 strains, and not all strains contained *Starships* (Gluck-Thaler et al., 2024). In *M. phaseolina*, four *Starships* and remnants were identified in 12 isolates (Gluck-Thaler et al., 2022). Overall, our study reveals an unprecedented abundance of *Starships* across *Verticillium* genomes.

We could unequivocally demonstrate that *Starships* make up about half of AGRs in the *V. dahliae* JR2 genome, while the remaining AGRs could not be assigned to *Starships*. AGRs were initially identified as lineage-specific genomic regions by comparisons among *V. dahliae* strains (de Jonge et al., 2013), while additional AGRs were subsequently identified based on their characteristic chromatin profile (Cook et al., 2020). Interestingly, the *Starship* regions identified in this study correspond to the initially identified AGRs. However, besides their chromatin profile, like *Starships*, also the non-*Starship* AGRs are enriched in horizontally transferred genes and genomic rearrangements. Thus, we speculate that these AGRs may be derived from unidentified *Starships* that became disrupted by genomic rearrangements. In support of this hypothesis, *Verticillium* genomes likely still contain many unidentified *Starship* regions, given 211 YR genes unassigned to known *Starship* regions in the 56 genomes, the majority (130/211) of which occur in AGRs. Overall, we conclude that *Starships* are the major constituents of AGRs in *V. dahliae*.

Many filamentous plant pathogens carry plastic genomic regions similar to AGRs (Dong et al., 2015; Kramer et al., 2023; Raffaele and Kamoun, 2012; Seidl et al., 2016; Seidl and Thomma, 2017), but how these regions evolved remains elusive (Torres et al., 2020). Although several *Starships* have been identified in Pezizomycotina plant pathogens (Bucknell et al., 2024; Gluck-Thaler et al., 2022; Gourlie et al., 2022; Haridas et al., 2023; Hill et al., 2024a; Peck et al., 2024; Tralamazza et al., 2024), the association of these *Starships* with the typical plastic genomic regions has not been noted previously.

HGT has driven virulence evolution in diverse fungal pathogens (Fitzpatrick, 2012; Soanes and Richards, 2014). HGT between phylogenetically close fungi is thought to be mediated by anastomosis and horizontal chromosome transfer, while mechanisms of HGT between phylogenetically distant fungi remains elusive (Fitzpatrick, 2012; Soanes and Richards, 2014). We reveal horizontal *Starship* transfer between *V. dahliae* and *V. nonalfalfae* on the one hand, and distantly related *Fusarium* fungi on the other hand. In addition, *Starship* regions in *V. dahliae* are enriched in cargo elements horizontally transferred to/from fungi belonging to further diverse Pezizomycotina orders. Thus, while previous studies have shown *Starship*-mediated HGT between fungi within the same order (Bucknell et al., 2024; Bucknell and McDonald, 2023; Gourlie et al., 2022; O’Donnell et al., 2024; Peck et al., 2024; A. S. Urquhart et al., 2024; Urquhart et al., 2023, 2022), our study extends this observation by revealing that *Starships* mediated HGT between phylogenetically distant fungi of different orders. There are many reports of cross-order HGT between diverse Pezizomycotina pathogens, though the mechanism remains generally unknown (Campbell et al., 2012; Hiruma et al., 2023; Khaldi and Wolfe, 2011; Kobayashi et al., 2023; Patron et al., 2007; Qiu et al., 2016; Slot and Rokas, 2011). Further identification of *Starships* in diverse taxa may help to understand the extent to which *Starships* have mediated such HGT events.

Pathogens exploit diverse effectors to colonize hosts, but it often remains unclear where, when, and how such effectors evolved, given that they are often considered lineage-specific inventions (Fouché et al., 2018). Since pathogen effectors often lack conserved protein domains, it has been speculated that such effectors may have evolved *de novo* (Fouché et al., 2018). *De novo* genes are defined to originate, at least in part, from non-coding DNA sequences, but the underlying evolutionary mechanisms remain poorly understood (McLysaght and Hurst, 2016; Oss and Carvunis, 2019). We show that *Starship* activity mediated the emergence of a virulence effector gene from a non-coding cargo sequence by the fusion with a section of a conserved effector gene. Given the strong association of *Starships* with plastic genomic regions enriched in effector genes, further elucidation of *Starship*-mediated gene evolution can be fundamental to understanding how novel virulence genes evolve.

## METHODS

### Genome assemblies and annotations

Genome sequences were collected from public databases or generated in this study (Supplementary Tables 1, 13, and 14). Fungal genomic DNA was extracted and sequenced using Oxford Nanopore Technologies as previously described (Chavarro-Carrero et al., 2021). Sequencing reads were assembled with Canu version 2.2 (Koren et al., 2017) and genome assembly quality was evaluated with BUSCO version 5.7.0 with the datasets eukaryota_odb10 and glomerellales_odb10 (Manni et al., 2021; Simão et al., 2015). The genome-wide average nucleotide identity (ANI) was calculated with FastANI version 1.33 (Jain et al., 2018).

Genes and TEs in *V. dahliae* strain JR2 were identified previously (Faino et al., 2015; Torres et al., 2021). In all other genomes, repetitive sequences were soft masked with RepeatModeler version 2.0.5 (Flynn et al., 2020) and RepeatMasker version 4.1.5 (Smit, AFA, Hubley, R & Green, P., 2013), after which genes were predicted with BRAKER version 3.0.8 (Gabriel et al., 2024) with GeneMark-EP+ version 4.72_lic (Brůna et al., 2020) and AUGUSTUS version 3.0.8 (Keller et al., 2011) using reference fungal proteins in OrthoDB version 11 (Kuznetsov et al., 2023). Orthologous gene groups were identified by eggNOG-mapper version 2.1.12 (Cantalapiedra et al., 2021) using eggNOG database version 5.0.2 (Huerta-Cepas et al., 2019). TEs were further characterized with the TE annotation pipeline EDTA version 2.2.1 (Ou et al., 2019) with curated TEs of *V. dahliae* VdLs17 (Amyotte et al., 2012).

### Identification of *Starships* and *Starship* regions

*Starships* and tyrosine recombinase (YR) genes were identified with Starfish v1.0.0 according to the standard workflow (Gluck-Thaler and Vogan, 2024). First, captain candidate YR genes were identified *de novo* from the genomes using the Starfish gene finder module with MetaEuk version 6.a5d39d9 (Levy Karin et al., 2020) and HMMER version 3.3.2 (Eddy, 2011) with the Pezizomycotina YR database attached to Starfish. The YR genes were grouped into families as defined previously (Gluck-Thaler and Vogan, 2024) based on a similarity search with HMMER version 3.3.2 (Eddy, 2011). The YR genes were then grouped into naves by clustering of *Verticillium* YRs using MMseqs2 version 14.7e284 easy-cluster (Steinegger and Söding, 2017) with thresholds of 50% amino acid sequence identity and 25% coverage. Each *Verticillium* YR navis was named with two letters of the YR family followed by an identifier number. Then, *Starship* candidates were identified via multiple sequence alignments among the 56 *Verticillium* genomes or among the 283 *Fusarium* genomes that were selected based on a relatively low number of contig numbers, suggesting a relatively highly contiguous genome assembly (Supplementary Table 1 and 14), by the Starfish element finder module using BLAST version 2.12.0+ (Camacho et al., 2009) and MUMmer version 4.0.0rc1 (Marçais et al., 2018). After the initial screening by Starfish, confident *Starships* were curated by 1) removing *Starship* candidates that lack flanking synteny over 20 kb against orthologous sites without *Starships* through manual inspection with Starfish pairViz, 2) removing *Starship* candidates that contain serial N stretches in scaffolds, and 3) unifying redundantly detected *Starships* due to the presence of YR genes on both strands around the 5’ ends. The *Starships* were grouped into haplotypes based on the captain navis and nucleotide *k*-mer similarities detected by sourmash version 4.8.3 (Pierce et al., 2019) with Starfish sim with default *k*-mer size 510 (Gluck-Thaler et al., 2024; Gluck-Thaler and Vogan, 2024), followed by the clustering using mcl version 14-137 (Enright et al., 2002) with minimal similarity threshold 0.05. Each *Verticillium Starship* haplotype was named after the captain navis ID followed by a suffix. Genomic regions orthologous to *Starship* insertion sites across *Verticillium* strains (Extended Data Fig. 4b) were identified by the Starfish region finder module based on the presence of low copy number (up to 5) eggNOG orthologs.

*Starship* regions were annotated by two approaches. First, regions downstream of YRs were annotated as *Starship* regions using Starfish extend (Gluck-Thaler and Vogan, 2024) with default settings using BLAST version 2.12.0+ (Camacho et al., 2009) based on similarity against curated *Starships*. Second, other *Starship* regions were identified irrespective of the presence of YR genes based on synteny against curated *Starships* through the alignments with nucmer of MUMmer version 4.0.0rc1 with options maxmatch and minimal alignment length 10,000, followed by filtering using delta-filter with the threshold of >80% nucleotide identity (Marçais et al., 2018). *Starships* identified in this study (Supplementary Tables 3 and 15) were used as reference. The pairwise sequence alignments between each *Starship* and syntenic *Starship* regions in each genome (Fig. 4b) were also performed using nucmer with the same settings.

The genomic compartments of *V. dahliae* strains JR2 and VdLs17 were assigned based on the overlap of *Starships* and *Starship* regions with the previously assigned AGRs core genomic regions (Cook et al., 2020), and centromeres (Seidl et al., 2020). AGRs in the JR2 genome were identified by chromatin profiling (Cook et al., 2020), while AGRs in the other strains refer to regions syntenic to AGRs in JR2 and regions absent in JR2 (Torres et al., 2024). To identify AGRs in strains other than JR2, genome alignments were performed with nucmer of MUMmer version 4.0.0rc1 (Marçais et al., 2018), followed by filtering of alignments with >80% nucleotide identity over 1 kb. The “*Starship*” compartment refers to *Starships* and additional *Starship* regions, “other AGR” refers to AGRs that do not belong to *Starship* regions, “centromere” refers to previously identified centromeres, and “other core” refers to core genomic regions that do not belong to *Starship* regions or centromeres. Genes and TEs in the JR2 genome were assigned to genomic compartments based on the location of their midpoints. TE counts over 10 kb windows in each genomic compartment were calculated using bedtools v2.30.0 intersect (Quinlan and Hall, 2010). Compartments smaller than 10 kb were omitted from the TE density analysis.

### Analysis of genomic syntenies and variations

Genomic syntenies and variations were identified with MUMmer version 4.0.0rc1 (Marçais et al., 2018). Specifically, segmental duplications in the JR2 genome were identified by genome self-alignment using nucmer with options maxmatch, nosimplify, and minimal alignment length 10,000, followed by the filtering with delta-filter. Inter-genomic syntenies and variations were identified by pairwise genome alignments with nucmer with option minimal alignment length 10,000, followed by the filtering and analysis by dnadiff. The genome-wide alignment coverage was calculated by integrating all fragmented hits. Among the structural variations (SVs) shown in this study, insertion/deletion (INDEL) corresponds to “GAP” and “BRK” in MUMmer4, while translocation (TRA) corresponds to “JMP” and “SEQ”. SV frequency was calculated by dividing the SV counts by the aligned length for each genomic compartment or genome. SV fold-enrichment was calculated by dividing the frequency in each genomic compartment by the genome-wide frequency.

### Transcriptome, epigenome, and 3D genome analyses

Global gene expression in *V. dahliae* strain JR2 was analyzed using the previous RNA sequencing (RNA-Seq) read data (Shi-Kunne et al., 2019) obtained from NCBI Short Read Archive (SRA) under the accession numbers listed in Supplementary Table 19. RNA-Seq reads were filtered with fastp version 0.19.5 (S. Chen et al., 2018) and mapped to the unmasked JR2 genome with STAR version 2.7.10a (Dobin et al., 2013) with previously applied options (Cook et al., 2020). The mapped reads were counted for each gene and TE by TEtranscripts version 2.2.1 with options stranded reverse and mode multi for 1,000 iterations (Jin et al., 2015). Fold change (FC) of gene expression was analyzed by DESeq2 version 1.42.1 (Love et al., 2014). Transcripts per million (TPM) were calculated using the established formula (Li et al., 2010). Mean TPM values of three biological replicates for individual genes/TEs were used for the data visualization and statistics.

Global histone H3K27me3 modifications in *V. dahliae* strain JR2 were analyzed using the previous chromatin immunoprecipitation sequencing (ChIP-Seq) data (Cook et al., 2020) obtained from NCBI SRA under the accession numbers listed in Supplementary Table 19. ChIP-Seq reads were filtered with fastp version 0.19.5 (S. Chen et al., 2018) and mapped to the JR2 genome masked as previously described (Cook et al., 2020) using BWA version 0.7.17 with BWA-MEM algorithm (Li, 2013). The mapped reads were sorted and indexed with samtools version 1.10 (Danecek et al., 2021) and counted with featureCounts version 2.0.1 (Liao et al., 2014) for each genomic compartment over 10 kb windows. The normalized counts per million (CPM) were calculated using the TPM formula (Li et al., 2010) by replacing transcripts with bins. The mean normalized CPM values of two biological replicates for respective bins were used for the data visualization and statistics.

Long-range chromatin interactions in *V. dahliae* strain JR2 were previously identified through chromatin conformation capture (Hi-C) (Torres et al., 2024). The previously analyzed data were directly used for visualization with the *Starship* positions.

### Local similarity searches

Local similarity searches were performed with BLAST version 2.15.0+ or 2.16.0+ (Camacho et al., 2009). The nucleotide-to-nucleotide searches were performed with the blastn algorithm with options word size 11 and e-value 0.05. The protein-to-translated nucleotide searches were performed with the tblastn algorithm with e-value threshold 0.05. The query *Verticillium* gene and TE sequences were obtained from GenBank and FungiDB under the accession numbers listed in Supplementary Table 20, and genes including introns were used as queries. The sequences and metadata of other virulence-associated proteins were obtained from PHI-base version 4.17 (Urban et al., 2022). Pezizomycotina genomes were obtained from Whole Genome Shotgun (WGS) sequences in GenBank under the accession numbers listed in Supplementary Table 13. Fungal taxonomic data were obtained from National Center for Biotechnology Information (NCBI) Taxonomy database. The coordinates of *Av2* orthologs (Supplementary Table 21) were identified by blastn search of the Pezizomycotina genomes with *V. dahliae Av2* or the *Av2* ortholog of *Fusarium phyllophilum* (*FpAv2*) under the accession numbers in Supplementary Table 20, followed by the selection of hits with >70% identity and >80% coverage.

Alien index (AI) values were calculated by the formula AI = log((best e-value for ingroup) + e-200) -log((best e-value for outgroup) + e-200) (Gladyshev et al., 2008) using hits with >50% coverage against *Verticillium* gene/TE queries by the blastn search. The e-value for no hits was set to 1 for queries with hits in either ingroup or outgroup, whereas AI values for queries with no hits in both ingroup and outgroup were not determined. The 332 non-*Verticillium* genomes of the order Glomerellales were used as ingroup, while the 9,641 genomes of the remaining 74 orders of Pezizomycotina were used as outgroup (Supplementary Table 13).

### Phylogenetic analyses

*Verticillium* phylogeny was inferred with REALPHY version 1.13 (Bertels et al., 2014) with TREE-PUZZLE version 5.3.rc16 (Schmidt et al., 2002) based on pairwise whole genome alignments with Bowtie version 2.2.5 (Langmead and Salzberg, 2012) with the JR2 genome as a reference.

*Starship* phylogeny was inferred by *k*-mer comparisons with mashtree version 1.4.6 with default *k*-mer size of 21, accuracy option mindepth 0, and bootstrapping for 1,000 iterations (Katz et al., 2019).

For the phylogenetic analyses of *Av2*, nucleotide sequences of *Av2* orthologs were extracted from the Pezizomycotina genomes at the coordinates listed in Supplementary Table 21 using SeqKit v2.3.0 subseq (Shen et al., 2024). The nucleotide sequences of *Av2* were aligned by MAFFT version 7.526 with L-INS-i method (Katoh and Standley, 2013), followed by trimming with trimAl version 1.4.rev15 with option strict (Capella-Gutiérrez et al., 2009). After multiple sequence alignments, phylogeny was inferred by IQ-TREE v2.0.3 (Minh et al., 2020) with the best substitution model TIM2e+G4 suggested by ModelFinder (Kalyaanamoorthy et al., 2017) and the ultrafast bootstrap approximation for 1,000 iterations (Minh et al., 2013).

For the phylogenetic analysis of NLPs, putative *Verticillium* proteins that contain an NPP1 domain (Pfam accession PF05630.16) (Seidl and Ackerveken, 2019) were identified using HMMER version 3.3.2 (Eddy, 2011). Putative secretion signals were identified with SignalP 6.0 (Teufel et al., 2022) or SignalP 3.0 (Dyrløv Bendtsen et al., 2004). The deduced NLP amino acid sequences were aligned and trimmed as described for *Av2*. Phylogeny of NLPs was also inferred as described for *Av2* with the different best substitution model VT+F+I+G4. NLP orthologs were numbered by the monophyletic relationship with NLP1 to NLP9 of *V. dahliae* strain VdLs17 (Klosterman et al., 2011; Santhanam et al., 2013) in the GenBank RefSeq database under the accession numbers listed in Supplementary Table 20.

### Targeted deletion of *NLP6* from the *Verticillium dahliae* genome

To generate an *NLP6* deletion construct, flanking sequences of its coding sequence were amplified from genomic DNA of *V. dahliae* strain VdLs17 using primers pair VDAG_04834-UP-F (GGTCTTAAUGAGAGTCGCAGAGGACTGAG) with VDAG_04834-UP-R (GGCATTAAUGCGGTGAGCAATGATAGATAATTGAG) to amplify the upstream region, and primers pair VDAG_04834-DOWN-F (GGACTTAAUGAGACGACCGACTTTGGGAG) with VDAG_04834-DOWN-R (GGGTTTAAUGGCGGATGCCAAAGCGCTC) to amplify the downstream region. The amplified products were cloned into pRF-HU2 as described previously (Frandsen et al., 2008), and subsequent *Agrobacterium tumefaciens*-mediated transformation of *V. dahliae* was performed as described previously (Santhanam, 2012). Transformants were selected on PDA supplemented with cefotaxime (Duchefa, Haarlem, The Netherlands) at 200 μg/ml and hygromycin (Duchefa) at 50 μg/ml, and homologous recombination was PCR-verified.

Pathogenicity assays were performed on ten-day-old tomato seedlings (MoneyMaker) plants using root-dip inoculation as previously described (Fradin et al., 2009). Disease symptoms were scored up to 14 dpi, pictures were taken, and ImageJ was used to determine canopy areas while fungal colonization was determined with real-time PCR. To this end, stem sections were taken at the height of the first internode, flash-frozen in liquid nitrogen, ground to powder, and genomic DNA was isolated. Real-time PCR was performed with a quantitative PCR core kit for SYBR Green I (Eurogentec, Seraing, Belgium) on an ABI7300 PCR machine (Applied Biosystems, Foster City, CA, U.S.A.). The *V. dahliae* internal transcribed spacer (ITS) levels were used relative to tomato ribulose-1,5-bisphosphate carboxylase/oxygenase (RuBisCo) levels to quantify fungal colonization of tomato plants (Snelders et al., 2023).

### Statistics

The normality and homoscedasticity of data were tested by Shapiro-Wilk test and Bartlett’s test, respectively, with the R base package ‘stats’ version 4.3.1 (R Core Team, 2023). The multiple comparisons of data with non-normal distributions and unequal variances (*p* < 0.05) were performed by Dunn’s test with the R package ‘dunn.test’ version 1.3.6 (Alexis Dinno, 2014). The multiple comparisons of data with normal distributions and equal variances (*p* < 0.05) were performed by Tukey’s test with the R package ‘multcomp’ version 1.4.26 (Hothorn et al., 2008).

### Data visualization

Circular and linear plots of genomic regions were generated with circlize version 0.4.16 (Gu et al., 2014) and gggenomes version 1.0.0 (Hackl et al., 2024), respectively. Phylogenetic trees were visualized with ggtree v3.10.1 (Yu et al., 2017). Other plots were generated with ggplot2 version 3.5.1 (Wickham, 2016).

## ACKNOWLEDGEMENTS

Y.S. acknowledges funding through an Overseas Research Fellowship from the Japan Society for the Promotion of Science. B.P.H.J.T. acknowledges funding by the Alexander von Humboldt Foundation in the framework of an Alexander von Humboldt Professorship endowed by the German Federal Ministry of Education and is furthermore supported by the Deutsche Forschungsgemeinschaft (DFG, German Research Foundation) under Germany’s Excellence Strategy – EXC 2048/1 – Project ID: 390686111. R.B. and M.H. acknowledge funding by the Research Foundation Flanders (FWO project ID: G004022N). We thank Dr. Edgar A. Chavarro-Carrero for providing sequencing read data of the *V. dahliae* genomes assembled in this study.

## AUTHOR CONTRIBUTIONS

Y.S. and B.P.H.J.T. conceived the project. Y.S., M.F.S., and B.P.H.J.T. designed the analyses. R.B. and M.H. contributed to the discovery of *NLP6* origin. G.C.M.B. performed fungal gene disruption and pathogenicity assays. Y.S., G.C.M.B, and B.P.H.J.T. analyzed the data. Y.S. and B.P.H.J.T. wrote the manuscript with feedback from all authors.

## EXTENDED DATA FIGURE LEGENDS

**Extended Data Fig. 1. Differential tyrosine recombinase (YR) repertoires across the *Verticillium* genus. a**, *Verticillium* phylogeny as described in Fig. 1a. **b**, Repertoires of YR naves per strain grouped by amino acid sequence similarity and named according to the YR family to which they belong. Heatmap colors and values indicate YR gene counts.

**Extended Data Fig. 2. Alignment of 35 *V. dahliae* genomes to the *V. dahliae* JR2 genome. a**, *Verticillium* phylogeny as described in Fig. 1a. **b**, Coverage plots of pairwise sequence alignments of 35 *V. dahliae* genomes to the JR2 genome in (**a**). Bar colors in JR2 represent the four genomic compartments “*Starship*”, “other AGR”, “centromere”, and “other core”. Bar colors for the 35 *V. dahliae* strains represent nucleotide identities of syntenic regions.

**Extended Data Fig. 3. *Verticillium Starship* cargo elements orthologous to virulence-associated genes of other fungi.** Hits (coverage and identity >80%) in the similarity search of *Verticillium Starships* with other fungal proteins that are described as virulence-associated in the Pathogen-Host Interactions Database (PHI-base) (Urban et al., 2022).

**Extended Data Fig. 4. *Starship* regions are dynamic. a**, *Verticillium* phylogeny as described in Fig. 1a. **b**, *Starship* occurrence in different genomic regions across the

*Verticillium* genomes. Columns indicate orthologous genomic regions with the presence/absence of *Starships* or of truncated *Starship*-like elements. **c-e**, Plasticity of *Starship* regions between *Verticillium* genomes including nested *Starship* insertion (**c**), *Starship* fragmentation (**d**), and invasion of different *Starships* (**e**). See Fig. 2a legend for the details of symbols.

**Extended Data Fig. 5. Search of Pezizomycotina genomes for *Verticillium Starship* orthologs. a**, Taxa of the input *Pezizomicotina* genomes used for the similarity search in Fig. 5. The left, middle, and right panels indicate the genomes of the phylum Pezizomycotina, the class Sordariomycetes, and the order Glomerellales, respectively. **b**, Coverage plots for *Verticillium Starships* with hits in the similarity search of Pezizomycotina genomes. Bar colors represent nucleotide identities for individual hits. Dashed red lines indicate the position of the G-LSR2 region (J.-Y. Chen et al., 2018).

**Extended Data Fig. 6. Horizontal transfer of *Av2* between *Verticillium* and *Fusarium*. a**, Nucleotide sequence alignment between *Av2* in *Verticillium dahliae* strain JR2 (*VdAv2*) and the ortholog in *Fusarium phyllophilum* strain NRRL 13617 (*FpAv2*). **b**, Phylogeny of *Av2* and its orthologs in *Verticillium* and *Fusarium*. Scale bars indicate nucleotide substitutions per site. Bootstrap values (>95%) for 1,000 iterations are shown at the nodes. **c**, Location of *Av2* in *Verticillium* and *Fusarium* chromosomes (chr), scaffolds (scf), or contigs (ctg). See Fig. 2a legend for the details of symbols. Ribbons connect syntenic regions between *Verticillium* and *Fusarium* genomic regions. Accession numbers of *Fusarium* genomes are described in Supplementary Table 21.

**Extended Data Fig. 7. Occurrence of active TEs in *Starships* of *Verticillium* and *Fusarium*. a**, Occurrence of transcriptionally active TEs characterized in *V. dahliae* strain VdLs17 (Amyotte et al., 2012) in diverse *Verticillium Starships* and adaptive genomic regions (AGRs). Circles in the left matrix represent the presence of the TEs (nucleotide identity >99% and coverage 100%) in the respective *Verticillium Starships* and VdLs17 genomic compartments. The circular plot shows the position of individual TEs with long triangles in/around *Starships* and *Starship* regions of *V. dahliae* strains JR2 and VdLs17. **b**, Occurrence of orthologs (nucleotide identity >70% and coverage >50%) of *Verticillium Starship* cargo TEs (**a**) in *Fusarium Starships*. Circles in the left matrix represent the presence of the respective TE orthologs in individual *Fusarium Starships*, with sizes representing TE coverage and colors representing nucleotide identity. The circular plot shows the position of individual TEs with long triangles in/around a *Starship* in *F. oxysporum* and *Starship* regions in VdLs17.

**Extended Data Fig. 8. Origin and virulence contribution of *NLP6* in *V. dahliae*. a**, Locations of *NLP3* and *NLP6* in *V. dahliae* strains JR2 and VdLs17. See Fig. 2a legend for the details of symbols. **b**, Amino acid sequence alignment between *NLP6* in *Verticillium dahliae* strain VdLs17 and NLP3 in *V. nubilum* strain 397 (*VnuNLP3*). The putative secretion signal and the conserved hexapeptide motif (Seidl and Ackerveken, 2019) are highlighted. **c**, Coverage plot based on the pairwise nucleotide sequence alignment between *NLP6* and *VnuNLP3*. **d-e**, Canopy area (**d**) and *V. dahliae* biomass (**e**) in tomato plants inoculated with wild-type (WT) and three independent *NLP6* deletion (Δ) lines of *V. dahliae* strain VdLs17 at 14 days post inoculation. Points indicate relative values for individual plants (*n* = 6) (**d**) or pooled plant samples (**e**), divided by the mean of WT-inoculated plants. Crossbars and error bars indicate mean ± standard deviation. Different letter labels indicate significant differences (Tukey’s test, *p* < 0.05).

## REFERENCES

Alexis Dinno, 2014. dunn.test: Dunn’s Test of Multiple Comparisons Using Rank Sums. 10.32614/CRAN.package.dunn.test

Amyotte, S.G., Tan, X., Pennerman, K. del Mar Jimenez-Gasco, M., Klosterman, S.J., Ma, L.-J., Dobinson, K.F., Veronese, P., 2012. Transposable elements in phytopathogenic Verticillium spp.: insights into genome evolution and inter-and intra-specific diversification. BMC Genomics 13, 314. 10.1186/1471-2164-13-314

Arkhipova, I.R., Yushenova, I.A., 2019. Giant Transposons in Eukaryotes: Is Bigger Better? Genome Biology and Evolution 11, 906–918. 10.1093/gbe/evz041

Bertels, F., Silander, O.K., Pachkov, M., Rainey, P.B., van Nimwegen, E., 2014. Automated Reconstruction of Whole-Genome Phylogenies from Short-Sequence Reads. Molecular Biology and Evolution 31, 1077–1088. 10.1093/molbev/msu088

Brůna, T., Lomsadze, A., Borodovsky, M., 2020. GeneMark-EP+: eukaryotic gene prediction with self-training in the space of genes and proteins. NAR Genomics and Bioinformatics 2, lqaa026. 10.1093/nargab/lqaa026

Bucknell, A., Wilson, H.M., Santos, K.C.G. do, Simpfendorfer, S., Milgate, A., Germain, H., Solomon, P.S., Bentham, A., McDonald, M.C., 2024. Sanctuary: A Starship transposon facilitating the movement of the virulence factor ToxA in fungal wheat pathogens. 10.1101/2024.03.04.583430

Bucknell, A.H., McDonald, M.C., 2023. That’s no moon, it’s a *Starship*: Giant transposons driving fungal horizontal gene transfer. Molecular Microbiology 120, 555–563. 10.1111/mmi.15118

Camacho, C., Coulouris, G., Avagyan, V., Ma, N., Papadopoulos, J., Bealer, K., Madden, T.L., 2009. BLAST+: architecture and applications. BMC Bioinformatics 10, 421. 10.1186/1471-2105-10-421

Campbell, M.A., Rokas, A., Slot, J.C., 2012. Horizontal Transfer and Death of a Fungal Secondary Metabolic Gene Cluster. Genome Biology and Evolution 4, 289–293. 10.1093/gbe/evs011

Cantalapiedra, C.P., Hernández-Plaza, A., Letunic, I., Bork, P., Huerta-Cepas, J., 2021. eggNOG-mapper v2: Functional Annotation, Orthology Assignments, and Domain Prediction at the Metagenomic Scale. Molecular Biology and Evolution 38, 5825–5829. 10.1093/molbev/msab293

Capella-Gutiérrez, S., Silla-Martínez, J.M., Gabaldón, T., 2009. trimAl: a tool for automated alignment trimming in large-scale phylogenetic analyses. Bioinformatics 25, 1972–1973. 10.1093/bioinformatics/btp348

Caracuel, Z., Roncero, M.I.G., Espeso, E.A., González-Verdejo, C.I., García-Maceira, F.I., Di Pietro, A., 2003. The pH signalling transcription factor PacC controls virulence in the plant pathogen *Fusarium oxysporum*. Molecular Microbiology 48, 765–779. 10.1046/j.1365-2958.2003.03465.x

Chavarro-Carrero, E.A., Vermeulen, J.P., E. Torres, D., Usami, T., Schouten, H.J., Bai, Y., Seidl, M.F., Thomma, B.P.H.J., 2021. Comparative genomics reveals the *in planta*-secreted *Verticillium dahliae* Av2 effector protein recognized in tomato plants that carry the *V2* resistance locus. Environmental Microbiology 23, 1941–1958. 10.1111/1462-2920.15288

Chen, J.-Y., Liu, C., Gui, Y.-J., Si, K.-W., Zhang, D.-D., Wang, J., Short, D.P.G., Huang, J.-Q., Li, N.-Y., Liang, Y., Zhang, W.-Q., Yang, L., Ma, X.-F., Li, T.-G., Zhou, L., Wang, B.-L., Bao, Y.-M., Subbarao, K.V., Zhang, G.-Y., Dai, X.-F., 2018. Comparative genomics reveals cotton-specific virulence factors in flexible genomic regions in *Verticillium dahliae* and evidence of horizontal gene transfer from *Fusarium*. New Phytologist 217, 756–770. 10.1111/nph.14861

Chen, S., Zhou, Y., Chen, Y., Gu, J., 2018. fastp: an ultra-fast all-in-one FASTQ preprocessor. Bioinformatics 34, i884–i890. 10.1093/bioinformatics/bty560

Cook, D.E., Kramer, H.M., Torres, D.E., Seidl, M.F., Thomma, B.P.H.J., 2020. A unique chromatin profile defines adaptive genomic regions in a fungal plant pathogen. eLife 9, e62208. 10.7554/eLife.62208

Cook, D.E., Mesarich, C.H., Thomma, B.P.H.J., 2015. Understanding Plant Immunity as a Surveillance System to Detect Invasion. Annual Review of Phytopathology 53, 541–563. 10.1146/annurev-phyto-080614-120114

Danecek, P., Bonfield, J.K., Liddle, J., Marshall, J., Ohan, V., Pollard, M.O., Whitwham, A., Keane, T., McCarthy, S.A., Davies, R.M., Li, H., 2021. Twelve years of SAMtools and BCFtools. GigaScience 10, giab008. 10.1093/gigascience/giab008

Dangl, J.L., Jones, J.D.G., 2024. A common immune response node in diverse plants. Science 386, 1344–1346. 10.1126/science.adu4930

de Jonge, R., Bolton, M.D., Kombrink, A., Berg, G.C.M. van den, Yadeta, K.A., Thomma, B.P.H.J., 2013. Extensive chromosomal reshuffling drives evolution of virulence in an asexual pathogen. Genome Res. 23, 1271–1282. 10.1101/gr.152660.112

de Jonge, R., Peter van Esse, H., Maruthachalam, K., Bolton, M.D., Santhanam, P., Saber, M.K., Zhang, Z., Usami, T., Lievens, B., Subbarao, K.V., Thomma, B.P.H.J. 2012. Tomato immune receptor Ve1 recognizes effector of multiple fungal pathogens uncovered by genome and RNA sequencing. Proceedings of the National Academy of Sciences 109, 5110–5115. 10.1073/pnas.1119623109

Depotter, J.R.L., Shi-Kunne, X., Missonnier, H., Liu, T., Faino, L., van den Berg, G.C.M., Wood, T.A., Zhang, B., Jacques, A., Seidl, M.F., Thomma, B.P.H.J., 2019. Dynamic virulence-related regions of the plant pathogenic fungus *Verticillium dahliae* display enhanced sequence conservation. Molecular Ecology 28, 3482–3495. 10.1111/mec.15168

Dobin, A., Davis, C.A., Schlesinger, F., Drenkow, J., Zaleski, C., Jha, S., Batut, P., Chaisson, M., Gingeras, T.R., 2013. STAR: ultrafast universal RNA-seq aligner. Bioinformatics 29, 15–21. 10.1093/bioinformatics/bts635

Dong, S., Raffaele, S., Kamoun, S., 2015. The two-speed genomes of filamentous pathogens: waltz with plants. Current Opinion in Genetics & Development, Genomes and evolution 35, 57–65. 10.1016/j.gde.2015.09.001

Dyrløv Bendtsen, J., Nielsen, H., von Heijne, G., Brunak, S., 2004. Improved Prediction of Signal Peptides: SignalP 3.0. Journal of Molecular Biology 340, 783–795. 10.1016/j.jmb.2004.05.028

Eddy, S.R., 2011. Accelerated Profile HMM Searches. PLOS Computational Biology 7, e1002195. 10.1371/journal.pcbi.1002195

Enright, A.J., Van Dongen, S., Ouzounis, C.A., 2002. An efficient algorithm for large-scale detection of protein families. Nucleic Acids Research 30, 1575–1584. 10.1093/nar/30.7.1575

Faino, L., Seidl, M.F., Datema, E., van den Berg, G.C.M., Janssen, A., Wittenberg, A.H.J., Thomma, B.P.H.J., 2015. Single-Molecule Real-Time Sequencing Combined with Optical Mapping Yields Completely Finished Fungal Genome. mBio 6, 10.1128/mbio.00936-15

Faino, L., Seidl, M.F., Shi-Kunne, X., Pauper, M., Berg, G.C.M. van den, Wittenberg, A.H.J., Thomma, B.P.H.J., 2016. Transposons passively and actively contribute to evolution of the two-speed genome of a fungal pathogen. Genome Res. 26, 1091–1100. 10.1101/gr.204974.116

Fedoroff, N.V., 2012. Transposable Elements, Epigenetics, and Genome Evolution. Science 338, 758–767. 10.1126/science.338.6108.758

Fitzpatrick, D.A., 2012. Horizontal gene transfer in fungi. FEMS Microbiology Letters 329, 1–8. 10.1111/j.1574-6968.2011.02465.x

Flynn, J.M., Hubley, R., Goubert, C., Rosen, J., Clark, A.G., Feschotte, C., Smit, A.F., 2020. RepeatModeler2 for automated genomic discovery of transposable element families. Proceedings of the National Academy of Sciences 117, 9451–9457. 10.1073/pnas.1921046117

Fouché, S., Plissonneau, C., Croll, D., 2018. The birth and death of effectors in rapidly evolving filamentous pathogen genomes. Current Opinion in Microbiology, Host Microbe Interactions: Fungi * Host Microbe Interactions: Parasitology 46, 34–42. 10.1016/j.mib.2018.01.020

Fradin, E.F., Thomma, B.P.H.J., 2006. Physiology and molecular aspects of *Verticillium* wilt diseases caused by *V. dahliae* and *V. albo-atrum*. Molecular Plant Pathology 7, 71–86. 10.1111/j.1364-3703.2006.00323.x

Fradin, E.F., Zhang, Z., Juarez Ayala, J.C., Castroverde, C.D.M., Nazar, R.N., Robb, J., Liu, C.-M., Thomma, B.P.H.J., 2009. Genetic Dissection of *Verticillium* Wilt Resistance Mediated by Tomato Ve1. Plant Physiology 150, 320–332. 10.1104/pp.109.136762

Frandsen, R.J., Andersson, J.A., Kristensen, M.B., Giese, H., 2008. Efficient four fragment cloning for the construction of vectors for targeted gene replacement in filamentous fungi. BMC Molecular Biology 9, 70. 10.1186/1471-2199-9-70

Frantzeskakis, L., Kusch, S., Panstruga, R., 2019. The need for speed: compartmentalized genome evolution in filamentous phytopathogens. Molecular Plant Pathology 20, 3–7. 10.1111/mpp.12738

Gabriel, L., Brůna, T., Hoff, K.J., Ebel, M., Lomsadze, A., Borodovsky, M., Stanke, M., 2024. BRAKER3: Fully automated genome annotation using RNA-seq and protein evidence with GeneMark-ETP, AUGUSTUS, and TSEBRA. Genome Res. 34, 769–777. 10.1101/gr.278090.123

Gladyshev, E.A., Meselson, M., Arkhipova, I.R., 2008. Massive Horizontal Gene Transfer in Bdelloid Rotifers. Science 320, 1210–1213. 10.1126/science.1156407

Gluck-Thaler, E., Forsythe, A., Puerner, C., Stajich, J.E., Croll, D., Cramer, R.A., Vogan, A.A., 2024. Giant transposons promote strain heterogeneity in a major fungal pathogen. 10.1101/2024.06.28.601215

Gluck-Thaler, E., Ralston, T., Konkel, Z., Ocampos, C.G., Ganeshan, V.D., Dorrance, A.E., Niblack, T.L., Wood, C.W., Slot, J.C., Lopez-Nicora, H.D., Vogan, A.A., 2022. Giant *Starship* Elements Mobilize Accessory Genes in Fungal Genomes. Molecular Biology and Evolution 39, msac109. 10.1093/molbev/msac109

Gluck-Thaler, E., Vogan, A.A., 2024. Systematic identification of cargo-mobilizing genetic elements reveals new dimensions of eukaryotic diversity. Nucleic Acids Research 52, 5496–5513. 10.1093/nar/gkae327

Gourlie, R., McDonald, M., Hafez, M., Ortega-Polo, R., Low, K.E., Abbott, D.W., Strelkov, S.E., Daayf, F., Aboukhaddour, R., 2022. The pangenome of the wheat pathogen *Pyrenophora tritici-repentis* reveals novel transposons associated with necrotrophic effectors *ToxA* and *ToxB*. BMC Biol 20, 239. 10.1186/s12915-022-01433-w

Gu, Z., Gu, L., Eils, R., Schlesner, M., Brors, B., 2014. circlize implements and enhances circular visualization in R. Bioinformatics 30, 2811–2812. 10.1093/bioinformatics/btu393

Hackl, T., Ankenbrand, M. J. & Adrichem, B. van. gggenomes: A Grammar of Graphics for Comparative Genomics. (2024).

Haridas, S., González, J.B., Riley, R., Koriabine, M., Yan, M., Ng, V., Rightmyer, A., Grigoriev, I.V., Baker, S.E., Turgeon, B.G., 2023. T-Toxin Virulence Genes: Unconnected Dots in a Sea of Repeats. mBio 14, e00261–23. 10.1128/mbio.00261-23

Harting, R., Starke, J., Kusch, H., Pöggeler, S., Maurus, I., Schlüter, R., Landesfeind, M., Bulla, I., Nowrousian, M., de Jonge, R., Stahlhut, G., Hoff, K.J., Aßhauer, K.P., Thürmer, A., Stanke, M., Daniel, R., Morgenstern, B., Thomma, B.P.H.J., Kronstad, J.W., Braus-Stromeyer, S.A., Braus, G.H., 2021. A 20-kb lineage-specific genomic region tames virulence in pathogenic amphidiploid *Verticillium longisporum*. Molecular Plant Pathology 22, 939–953. 10.1111/mpp.13071

Hill, R., Grey, M., Fedi, M.O., Smith, D., Ward, S.J., Canning, G., Irish, N., Smith, J., McMillan, V.E., Hammond, J., Osborne, S.-J., Chancellor, T., Swarbreck, D., Hall, N., Palma-Guerrero, J., Hammond-Kosack, K.E., McMullan, M., 2024a. Evolutionary genomics reveals variation in structure and genetic content implicated in virulence and lifestyle in the genus Gaeumannomyces. 10.1101/2024.02.15.580261

Hill, R., Smith, D., Canning, G., Grey, M., Hammond-Kosack, K., McMullan, M., 2024b. Starship giant transposable elements cluster by host taxonomy using kmer-based phylogenetics. 10.1101/2024.08.30.610507

Hiruma, K., Aoki, S., Takino, J., Higa, T., Utami, Y.D., Shiina, A., Okamoto, M., Nakamura, M., Kawamura, N., Ohmori, Y., Sugita, R., Tanoi, K., Sato, T., Oikawa, H., Minami, A., Iwasaki, W., Saijo, Y., 2023. A fungal sesquiterpene biosynthesis gene cluster critical for mutualist-pathogen transition in *Colletotrichum tofieldiae*. Nat Commun 14, 5288. 10.1038/s41467-023-40867-w

Hothorn, T., Bretz, F., Westfall, P., 2008. Simultaneous Inference in General Parametric Models. Biometrical Journal 50, 346–363. 10.1002/bimj.200810425

Huerta-Cepas, J., Szklarczyk, D., Heller, D., Hernández-Plaza, A., Forslund, S.K., Cook, H., Mende, D.R., Letunic, I., Rattei, T., Jensen, L.J., von Mering, C., Bork, P., 2019. eggNOG 5.0: a hierarchical, functionally and phylogenetically annotated orthology resource based on 5090 organisms and 2502 viruses. Nucleic Acids Research 47, D309–D314. 10.1093/nar/gky1085

Inderbitzin, P., Bostock, R.M., Davis, R.M., Usami, T., Platt, H.W., Subbarao, K.V., 2011. Phylogenetics and Taxonomy of the Fungal Vascular Wilt Pathogen *Verticillium*, with the Descriptions of Five New Species. PLOS ONE 6, e28341. 10.1371/journal.pone.0028341

Inoue, Y., Takeda, H., 2023. Teratorn and its relatives – a cross-point of distinct mobile elements, transposons and viruses. Front. Vet. Sci. 10. 10.3389/fvets.2023.1158023

Jain, C., Rodriguez-R, L.M., Phillippy, A.M., Konstantinidis, K.T., Aluru, S., 2018. High throughput ANI analysis of 90K prokaryotic genomes reveals clear species boundaries. Nat Commun 9, 5114. 10.1038/s41467-018-07641-9

Jin, Y., Tam, O.H., Paniagua, E., Hammell, M., 2015. TEtranscripts: a package for including transposable elements in differential expression analysis of RNA-seq datasets. Bioinformatics 31, 3593–3599. 10.1093/bioinformatics/btv422

Kalyaanamoorthy, S., Minh, B.Q., Wong, T.K.F., von Haeseler, A., Jermiin, L.S., 2017. ModelFinder: fast model selection for accurate phylogenetic estimates. Nat Methods 14, 587–589. 10.1038/nmeth.4285

Katoh, K., Standley, D.M., 2013. MAFFT Multiple Sequence Alignment Software Version 7: Improvements in Performance and Usability. Molecular Biology and Evolution 30, 772–780. 10.1093/molbev/mst010

Katz, L.S., Griswold, T., Morrison, S.S., Caravas, J.A., Zhang, S., Bakker, H.C. den, Deng, X., Carleton, H.A., 2019. Mashtree: a rapid comparison of whole genome sequence files. Journal of Open Source Software 4, 1762. 10.21105/joss.01762

Keller, O., Kollmar, M., Stanke, M., Waack, S., 2011. A novel hybrid gene prediction method employing protein multiple sequence alignments. Bioinformatics 27, 757–763. 10.1093/bioinformatics/btr010

Khaldi, N., Wolfe, K.H., 2011. Evolutionary Origins of the Fumonisin Secondary Metabolite Gene Cluster in *Fusarium verticillioides* and *Aspergillus niger*. Int J Evol Biol 2011, 423821. 10.4061/2011/423821

Klimes, A., Dobinson, K.F., Thomma, B.P.H.J., Klosterman, S.J., 2015. Genomics Spurs Rapid Advances in Our Understanding of the Biology of Vascular Wilt Pathogens in the Genus *Verticillium*. Annual Review of Phytopathology 53, 181–198. 10.1146/annurev-phyto-080614-120224

Klosterman, S.J., Subbarao, K.V., Kang, S., Veronese, P., Gold, S.E., Thomma, B.P.H.J., Chen, Z., Henrissat, B., Lee, Y.-H., Park, J., Garcia-Pedrajas, M.D., Barbara, D.J., Anchieta, A., Jonge, R. de Santhanam, P., Maruthachalam, K., Atallah, Z., Amyotte, S.G., Paz, Z., Inderbitzin, P., Hayes, R.J., Heiman, D.I., Young, S., Zeng, Q., Engels, R., Galagan, J., Cuomo, C.A., Dobinson, K.F., Ma, L.-J., 2011. Comparative Genomics Yields Insights into Niche Adaptation of Plant Vascular Wilt Pathogens. PLOS Pathogens 7, e1002137. 10.1371/journal.ppat.1002137

Kobayashi, N., Dang, T.A., Pham, K.T.M., Gómez Luciano, L.B., Van Vu, B., Izumitsu, K., Shimizu, M., Ikeda, K., Li, W.-H., Nakayashiki, H., 2023. Horizontally Transferred DNA in the Genome of the Fungus *Pyricularia oryzae* is Associated With Repressive Histone Modifications. Molecular Biology and Evolution 40, msad186. 10.1093/molbev/msad186

Kodama, S., Ishizuka, J., Miyashita, I., Ishii, T., Nishiuchi, T., Miyoshi, H., Kubo, Y., 2017. The morphogenesis-related NDR kinase pathway of *Colletotrichum orbiculare* is required for translating plant surface signals into infection-related morphogenesis and pathogenesis. PLOS Pathogens 13, e1006189. 10.1371/journal.ppat.1006189

Kombrink, A., Rovenich, H., Shi-Kunne, X., Rojas-Padilla, E., van den Berg, G.C.M., Domazakis, E., de Jonge, R., Valkenburg, D.-J., Sánchez-Vallet, A., Seidl, M.F., Thomma, B.P.H.J., 2017. *Verticillium dahliae* LysM effectors differentially contribute to virulence on plant hosts. Molecular Plant Pathology 18, 596–608. 10.1111/mpp.12520

Koren, S., Walenz, B.P., Berlin, K., Miller, J.R., Bergman, N.H., Phillippy, A.M., 2017. Canu: scalable and accurate long-read assembly via adaptive *k*-mer weighting and repeat separation. Genome Res. 27, 722–736. 10.1101/gr.215087.116

Kramer, H.M., Cook, D.E., Seidl, M.F., Thomma, B.P.H.J., 2023. Epigenetic regulation of nuclear processes in fungal plant pathogens. PLOS Pathogens 19, e1011525. 10.1371/journal.ppat.1011525

Kramer, H.M., Cook, D.E., van den Berg, G.C.M., Seidl, M.F., Thomma, B.P.H.J., 2021. Three putative DNA methyltransferases of *Verticillium dahliae* differentially contribute to DNA methylation that is dispensable for growth, development and virulence. Epigenetics & Chromatin 14, 21. 10.1186/s13072-021-00396-6

Kramer, H.M., Seidl, M.F., Thomma, B.P.H.J., Cook, D.E., 2022. Local Rather than Global H3K27me3 Dynamics Are Associated with Differential Gene Expression in *Verticillium dahliae*. mBio 13, e03566–21. 10.1128/mbio.03566-21

Kuznetsov, D., Tegenfeldt, F., Manni, M., Seppey, M., Berkeley, M., Kriventseva, E.V., Zdobnov, E.M., 2023. OrthoDB v11: annotation of orthologs in the widest sampling of organismal diversity. Nucleic Acids Research 51, D445–D451. 10.1093/nar/gkac998

Langmead, B., Salzberg, S.L., 2012. Fast gapped-read alignment with Bowtie 2. Nat Methods 9, 357–359. 10.1038/nmeth.1923

Levy Karin, E., Mirdita, M., Söding, J., 2020. MetaEuk—sensitive, high-throughput gene discovery, and annotation for large-scale eukaryotic metagenomics. Microbiome 8, 48. 10.1186/s40168-020-00808-x

Li, B., Ruotti, V., Stewart, R.M., Thomson, J.A., Dewey, C.N., 2010. RNA-Seq gene expression estimation with read mapping uncertainty. Bioinformatics 26, 493–500. 10.1093/bioinformatics/btp692

Li, H., 2013. Aligning sequence reads, clone sequences and assembly contigs with BWA-MEM. 10.48550/arXiv.1303.3997

Liao, Y., Smyth, G.K., Shi, W., 2014. featureCounts: an efficient general purpose program for assigning sequence reads to genomic features. Bioinformatics 30, 923–930. 10.1093/bioinformatics/btt656

Love, M.I., Huber, W., Anders, S., 2014. Moderated estimation of fold change and dispersion for RNA-seq data with DESeq2. Genome Biology 15, 550. 10.1186/s13059-014-0550-8

Manni, M., Berkeley, M.R., Seppey, M., Simão, F.A., Zdobnov, E.M., 2021. BUSCO Update: Novel and Streamlined Workflows along with Broader and Deeper Phylogenetic Coverage for Scoring of Eukaryotic, Prokaryotic, and Viral Genomes. Molecular Biology and Evolution 38, 4647–4654. 10.1093/molbev/msab199

Marçais, G., Delcher, A.L., Phillippy, A.M., Coston, R., Salzberg, S.L., Zimin, A., 2018. MUMmer4: A fast and versatile genome alignment system. PLOS Computational Biology 14, e1005944. 10.1371/journal.pcbi.1005944

McLysaght, A., Hurst, L.D., 2016. Open questions in the study of *de novo* genes: what, how and why. Nat Rev Genet 17, 567–578. 10.1038/nrg.2016.78

Minh, B.Q., Nguyen, M.A.T., von Haeseler, A., 2013. Ultrafast Approximation for Phylogenetic Bootstrap. Molecular Biology and Evolution 30, 1188–1195. 10.1093/molbev/mst024

Minh, B.Q., Schmidt, H.A., Chernomor, O., Schrempf, D., Woodhams, M.D., von Haeseler, A., Lanfear, R., 2020. IQ-TREE 2: New Models and Efficient Methods for Phylogenetic Inference in the Genomic Era. Molecular Biology and Evolution 37, 1530–1534. 10.1093/molbev/msaa015

Möller, M., Stukenbrock, E.H., 2017. Evolution and genome architecture in fungal plant pathogens. Nat Rev Microbiol 15, 756–771. 10.1038/nrmicro.2017.76

Nguyen, Q.B., Kadotani, N., Kasahara, S., Tosa, Y., Mayama, S., Nakayashiki, H., 2008. Systematic functional analysis of calcium-signalling proteins in the genome of the rice-blast fungus, *Magnaporthe oryzae*, using a high-throughput RNA-silencing system. Molecular Microbiology 68, 1348–1365. 10.1111/j.1365-2958.2008.06242.x

O’Donnell, S., Rezende, G., Vernadet, J.-P., Snirc, A., Ropars, J., 2024. Harbouring Starships: The accumulation of large Horizontal Gene Transfers in Domesticated and Pathogenic Fungi. 10.1101/2024.07.03.601904

Ondov, B.D., Treangen, T.J., Melsted, P., Mallonee, A.B., Bergman, N.H., Koren, S., Phillippy, A.M., 2016. Mash: fast genome and metagenome distance estimation using MinHash. Genome Biology 17, 132. 10.1186/s13059-016-0997-x

Ortoneda, M., Guarro, J., Madrid, M.P., Caracuel, Z., Roncero, M.I.G., Mayayo, E., Di Pietro, A., 2004. *Fusarium oxysporum* as a Multihost Model for the Genetic Dissection of Fungal Virulence in Plants and Mammals. Infection and Immunity 72, 1760–1766. 10.1128/iai.72.3.1760-1766.2004

Oss, S.B.V., Carvunis, A.-R., 2019. *De novo* gene birth. PLOS Genetics 15, e1008160. 10.1371/journal.pgen.1008160

Ou, S., Su, W., Liao, Y., Chougule, K., Agda, J.R.A., Hellinga, A.J., Lugo, C.S.B., Elliott, T.A., Ware, D., Peterson, T., Jiang, N., Hirsch, C.N., Hufford, M.B., 2019. Benchmarking transposable element annotation methods for creation of a streamlined, comprehensive pipeline. Genome Biology 20, 275. 10.1186/s13059-019-1905-y

Patron, N.J., Waller, R.F., Cozijnsen, A.J., Straney, D.C., Gardiner, D.M., Nierman, W.C., Howlett, B.J., 2007. Origin and distribution of epipolythiodioxopiperazine (ETP) gene clusters in filamentous ascomycetes. BMC Evolutionary Biology 7, 174. 10.1186/1471-2148-7-174

Peck, L.D., Llewellyn, T., Bennetot, B., O’Donnell, S., Nowell, R.W., Ryan, M.J., Flood, J., Vega, R.C.R. de la, Ropars, J., Giraud, T., Spanu, P.D., Barraclough, T.G., 2024. Horizontal transfers between fungal *Fusarium* species contributed to successive outbreaks of coffee wilt disease. PLOS Biology 22, e3002480. 10.1371/journal.pbio.3002480

Pierce, N.T., Irber, L., Reiter, T., Brooks, P., Brown, C.T., 2019. Large-scale sequence comparisons with sourmash. 10.12688/f1000research.19675.1

Qiu, H., Cai, G., Luo, J., Bhattacharya, D., Zhang, N., 2016. Extensive horizontal gene transfers between plant pathogenic fungi. BMC Biology 14, 41. 10.1186/s12915-016-0264-3

Quinlan, A.R., Hall, I.M., 2010. BEDTools: a flexible suite of utilities for comparing genomic features. Bioinformatics 26, 841–842. 10.1093/bioinformatics/btq033

R Core Team, 2023. R: A Language and Environment for Statistical Computing. R Foundation for Statistical Computing, Vienna, Austria.

Raffaele, S., Kamoun, S., 2012. Genome evolution in filamentous plant pathogens: why bigger can be better. Nat Rev Microbiol 10, 417–430. 10.1038/nrmicro2790

Ravenhall, M., Škunca, N., Lassalle, F., Dessimoz, C., 2015. Inferring Horizontal Gene Transfer. PLOS Computational Biology 11, e1004095. 10.1371/journal.pcbi.1004095

Santhanam, P., 2012. Random Insertional Mutagenesis in Fungal Genomes to Identify Virulence Factors, in: Bolton, M.D., Thomma, B.P.H.J. (Eds.), Plant Fungal Pathogens: Methods and Protocols. Humana Press, Totowa, NJ, pp. 509–517. 10.1007/978-1-61779-501-5_31

Santhanam, P., van Esse, H.P., Albert, I., Faino, L., Nürnberger, T., Thomma, B.P.H.J., 2013. Evidence for Functional Diversification Within a Fungal NEP1-Like Protein Family. MPMI 26, 278–286. 10.1094/MPMI-09-12-0222-R

Schmidt, H.A., Strimmer, K., Vingron, M., von Haeseler, A., 2002. TREE-PUZZLE: maximum likelihood phylogenetic analysis using quartets and parallel computing. Bioinformatics 18, 502–504. 10.1093/bioinformatics/18.3.502

Seidl, M.F., Ackerveken, G.V. den, 2019. Activity and Phylogenetics of the Broadly Occurring Family of Microbial Nep1-Like Proteins. Annual Review of Phytopathology 57, 367–386. 10.1146/annurev-phyto-082718-100054

Seidl, M.F., Cook, D.E., Thomma, B.P.H.J., 2016. Chromatin Biology Impacts Adaptive Evolution of Filamentous Plant Pathogens. PLOS Pathogens 12, e1005920. 10.1371/journal.ppat.1005920

Seidl, M.F., Kramer, H.M., Cook, D.E., Fiorin, G.L., van den Berg, G.C.M., Faino, L., Thomma, B.P.H.J., 2020. Repetitive Elements Contribute to the Diversity and Evolution of Centromeres in the Fungal Genus *Verticillium*. mBio 11, 10.1128/mbio.01714-20.

Seidl, M.F., Thomma, B.P.H.J., 2017. Transposable Elements Direct The Coevolution between Plants and Microbes. Trends in Genetics, Transposable Elements 33, 842–851. 10.1016/j.tig.2017.07.003

Shen, W., Sipos, B., Zhao, L., 2024. SeqKit2: A Swiss army knife for sequence and alignment processing. iMeta 3, e191. 10.1002/imt2.191

Shi-Kunne, X., Faino, L., van den Berg, G.C.M., Thomma, B.P.H.J., Seidl, M.F., 2018. Evolution within the fungal genus *Verticillium* is characterized by chromosomal rearrangement and gene loss. Environmental Microbiology 20, 1362–1373. 10.1111/1462-2920.14037

Shi-Kunne, X., Jové, R. de P., Depotter, J.R.L., Ebert, M.K., Seidl, M.F., Thomma, B.P.H.J., 2019. *In silico* prediction and characterisation of secondary metabolite clusters in the plant pathogenic fungus *Verticillium dahliae*. FEMS Microbiology Letters 366, fnz081. 10.1093/femsle/fnz081

Simão, F.A., Waterhouse, R.M., Ioannidis, P., Kriventseva, E.V., Zdobnov, E.M., 2015. BUSCO: assessing genome assembly and annotation completeness with single-copy orthologs. Bioinformatics 31, 3210–3212.10.1093/bioinformatics/btv351

Slot, J.C., Rokas, A., 2011. Horizontal Transfer of a Large and Highly Toxic Secondary Metabolic Gene Cluster between Fungi. Current Biology 21, 134–139. 10.1016/j.cub.2010.12.020

Smit, AFA, Hubley, R & Green, P., 2013. RepeatMasker Open-4.0.

Snelders, N.C., Boshoven, J.C., Song, Y., Schmitz, N., Fiorin, G.L., Rovenich, H., van den Berg, G.C.M., Torres, D.E., Petti, G.C., Prockl, Z., Faino, L., Seidl, M.F., Thomma, B.P.H.J., 2023. A highly polymorphic effector protein promotes fungal virulence through suppression of plant-associated Actinobacteria. New Phytologist 237, 944–958. 10.1111/nph.18576

Snelders, N.C., Rovenich, H., Petti, G.C., Rocafort, M., van den Berg, G.C.M., Vorholt, J.A., Mesters, J.R., Seidl, M.F., Nijland, R., Thomma, B.P.H.J., 2020. Microbiome manipulation by a soil-borne fungal plant pathogen using effector proteins. Nat. Plants 6, 1365–1374. 10.1038/s41477-020-00799-5

Soanes, D., Richards, T.A., 2014. Horizontal Gene Transfer in Eukaryotic Plant Pathogens. Annual Review of Phytopathology 52, 583–614. 10.1146/annurev-phyto-102313-050127

Steinegger, M., Söding, J., 2017. MMseqs2 enables sensitive protein sequence searching for the analysis of massive data sets. Nat Biotechnol 35, 1026–1028. 10.1038/nbt.3988

Teufel, F., Almagro Armenteros, J.J., Johansen, A.R., Gíslason, M.H., Pihl, S.I., Tsirigos, K.D., Winther, O., Brunak, S., von Heijne, G., Nielsen, H., 2022. SignalP 6.0 predicts all five types of signal peptides using protein language models. Nat Biotechnol 40, 1023–1025. 10.1038/s41587-021-01156-3

Torres, D.E., Kramer, H.M., Tracanna, V., Fiorin, G.L., Cook, D.E., Seidl, M.F., Thomma, B.P.H.J., 2024. Implications of the three-dimensional chromatin organization for genome evolution in a fungal plant pathogen. Nat Commun 15, 1701. 10.1038/s41467-024-45884-x

Torres, D.E., Oggenfuss, U., Croll, D., Seidl, M.F., 2020. Genome evolution in fungal plant pathogens: looking beyond the two-speed genome model. Fungal Biology Reviews 34, 136–143. 10.1016/j.fbr.2020.07.001

Torres, D.E., Thomma, B.P.H.J., Seidl, M.F., 2021. Transposable Elements Contribute to Genome Dynamics and Gene Expression Variation in the Fungal Plant Pathogen Verticillium dahliae. Genome Biology and Evolution 13, evab135. 10.1093/gbe/evab135

Tralamazza, S.M., Gluck-Thaler, E., Feurtey, A., Croll, D., 2024. Copy number variation introduced by a massive mobile element facilitates global thermal adaptation in a fungal wheat pathogen. Nat Commun 15, 5728. 10.1038/s41467-024-49913-7

Urban, M., Cuzick, A., Seager, J., Wood, V., Rutherford, K., Venkatesh, S.Y., Sahu, J., Iyer, S.V., Khamari, L., De Silva, N., Martinez, M.C., Pedro, H., Yates, A.D., Hammond-Kosack, K.E., 2022. PHI-base in 2022: a multi-species phenotype database for Pathogen–Host Interactions. Nucleic Acids Research 50, D837–D847. 10.1093/nar/gkab1037

Urquhart, A., Vogan, A.A., Gluck-Thaler, E., 2024. *Starships*: a new frontier for fungal biology. Trends in Genetics 40, 1060–1073. 10.1016/j.tig.2024.08.006

Urquhart, A.S., Chong, N.F., Yang, Y., Idnurm, A., 2022. A large transposable element mediates metal resistance in the fungus *Paecilomyces variotii*. Current Biology 32, 937–950.e5. 10.1016/j.cub.2021.12.048

Urquhart, A.S., Gluck-Thaler, E., Vogan, A.A., 2024. Gene acquisition by giant transposons primes eukaryotes for rapid evolution via horizontal gene transfer. Science Advances 10, eadp8738. 10.1126/sciadv.adp8738

Urquhart, A.S., Vogan, A.A., Gardiner, D.M., Idnurm, A., 2023. *Starships* are active eukaryotic transposable elements mobilized by a new family of tyrosine recombinases. Proceedings of the National Academy of Sciences 120, e2214521120. 10.1073/pnas.2214521120

Vogan, A.A., Ament-Velásquez, S.L., Bastiaans, E., Wallerman, O., Saupe, S.J., Suh, A., Johannesson, H., 2021. The *Enterprise*, a massive transposon carrying *Spok* meiotic drive genes. Genome Res. 31, 789–798. 10.1101/gr.267609.120

Wicker, T., Sabot, F., Hua-Van, A., Bennetzen, J.L., Capy, P., Chalhoub, B., Flavell, A., Leroy, P., Morgante, M., Panaud, O., Paux, E., SanMiguel, P., Schulman, A.H., 2007. A unified classification system for eukaryotic transposable elements. Nat Rev Genet 8, 973–982. 10.1038/nrg2165

Wickham, H. ggplot2: Elegant Graphics for Data Analysis. (Springer-Verlag New York, 2016).

Yu, G., Smith, D.K., Zhu, H., Guan, Y., Lam, T.T.-Y., 2017. ggtree: an r package for visualization and annotation of phylogenetic trees with their covariates and other associated data. Methods in Ecology and Evolution 8, 28–36. 10.1111/2041-210X.12628

Zhang, J., Jin, K., Xia, Y., 2017. Contributions of β-tubulin to cellular morphology, sporulation and virulence in the insect-fungal pathogen, *Metarhizium acridum*. Fungal Genetics and Biology 103, 16–24. 10.1016/j.fgb.2017.03.005

