## Supplementary figures and images for "*Starship* giant transposons dominate plastic genomic regions in a fungal plant pathogen and drive virulence evolution"

### Supplemental Figure 1

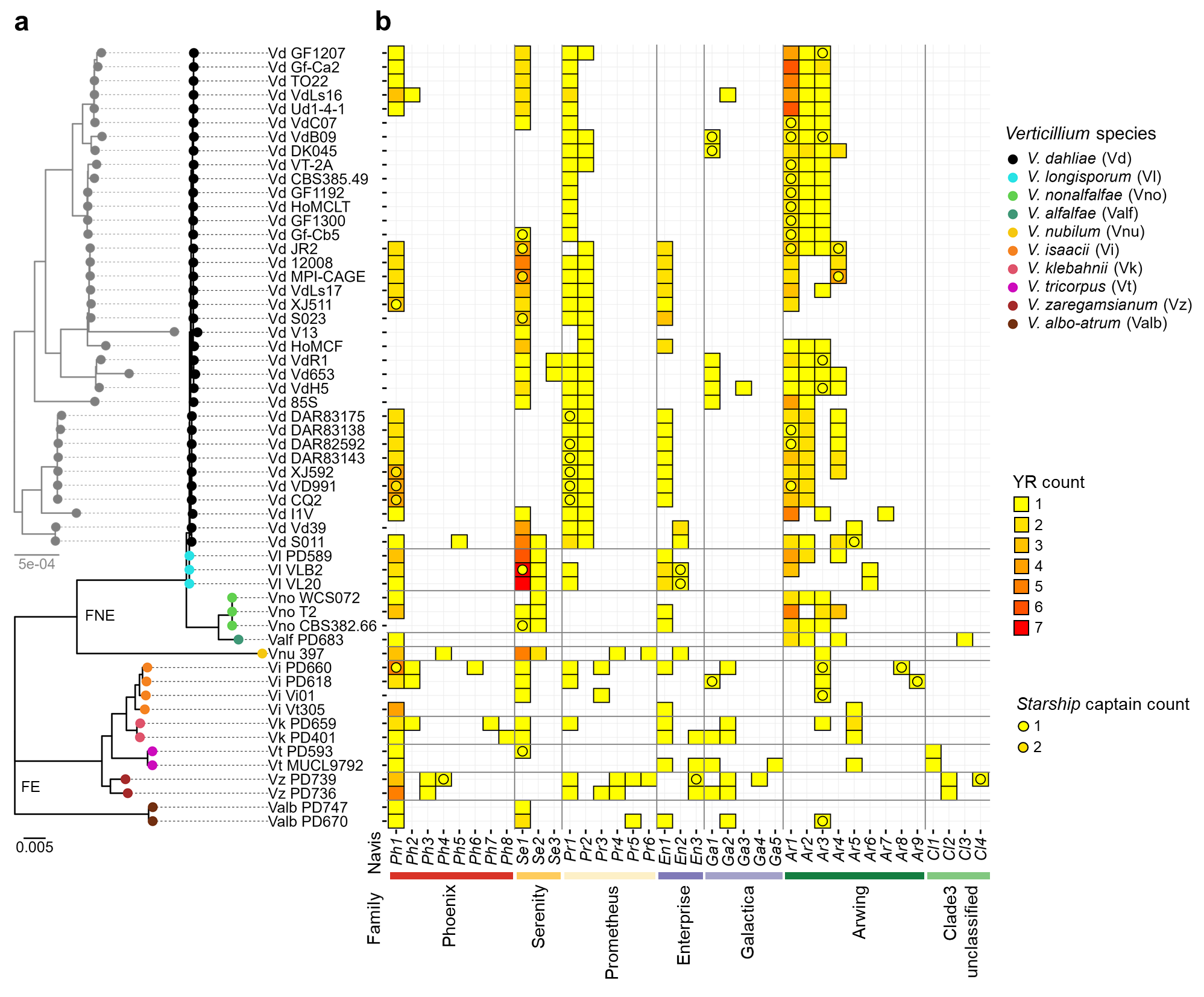

### Supplemental Figure 2

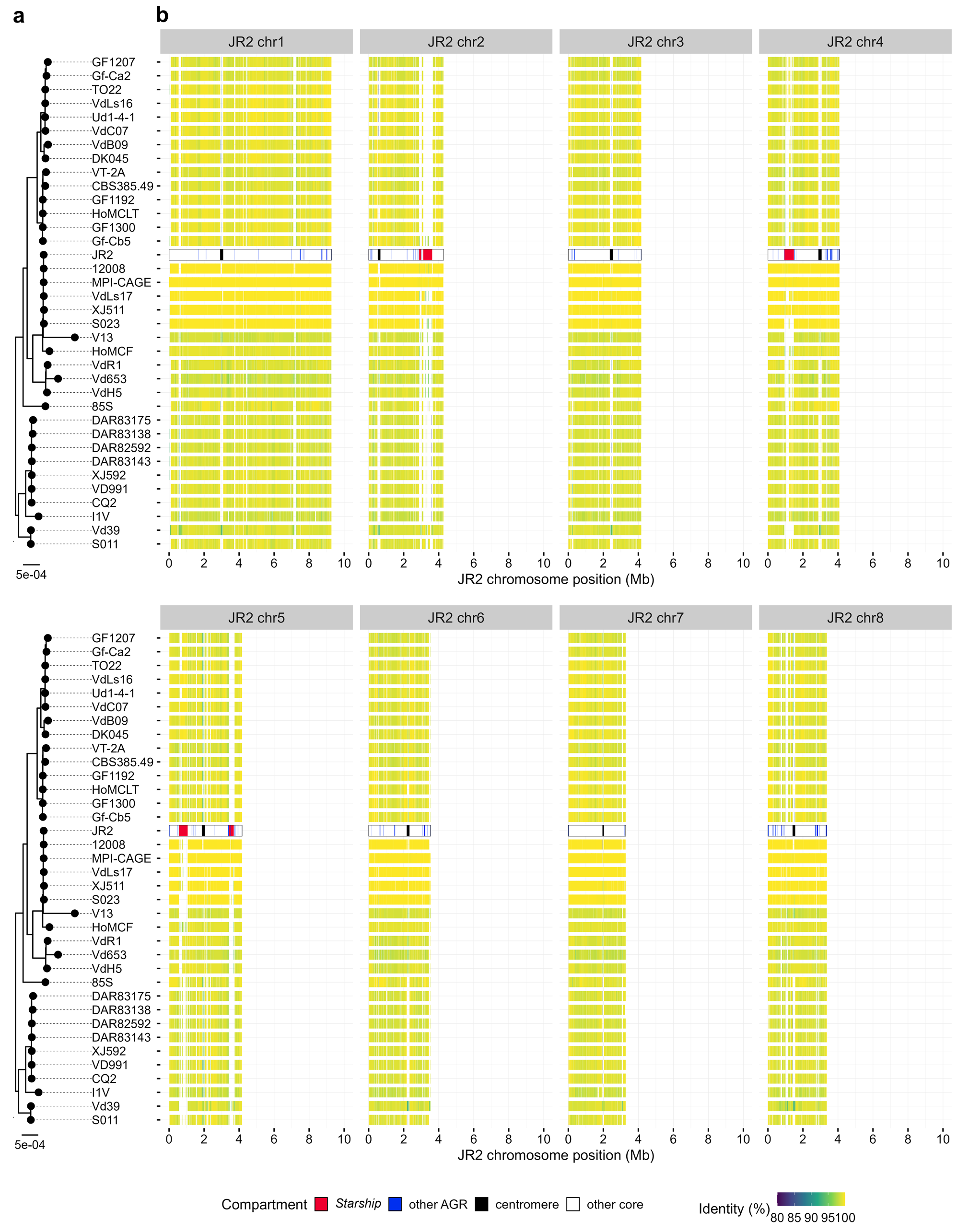

### Supplemental Figure 3

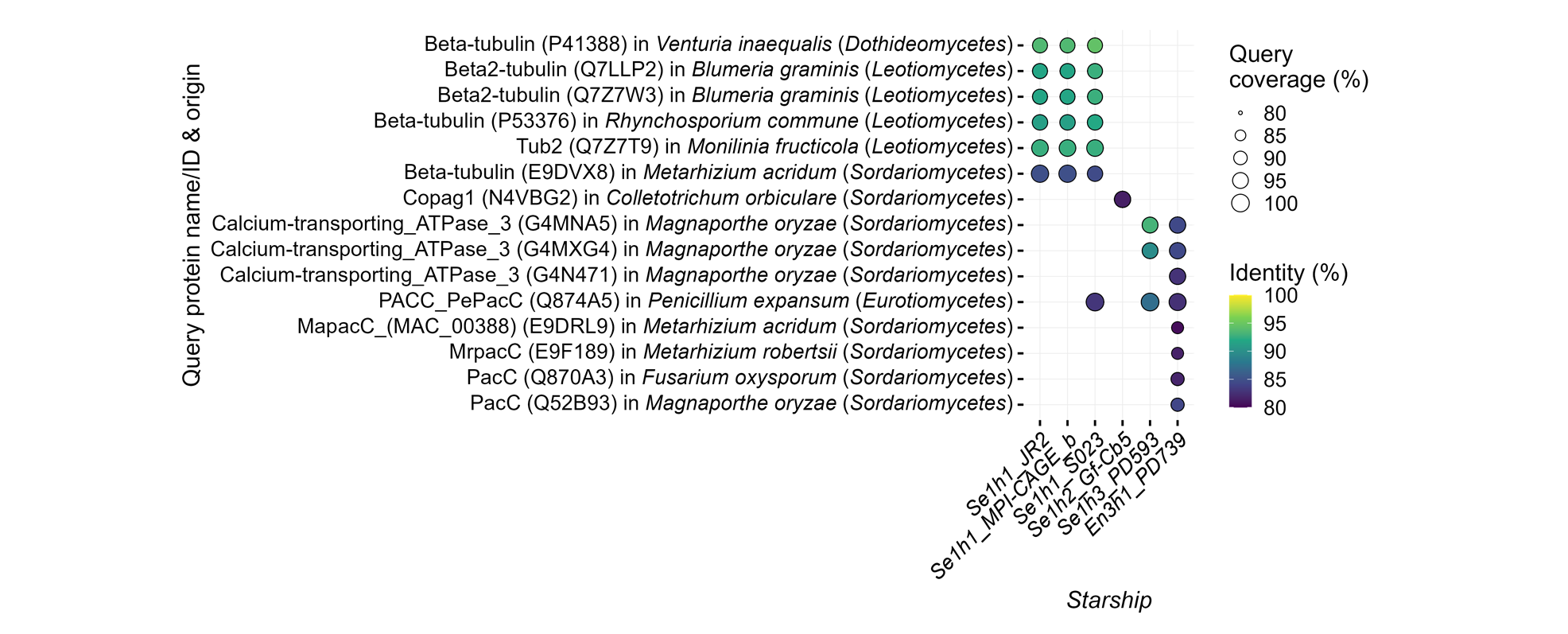

### Supplemental Figure 4

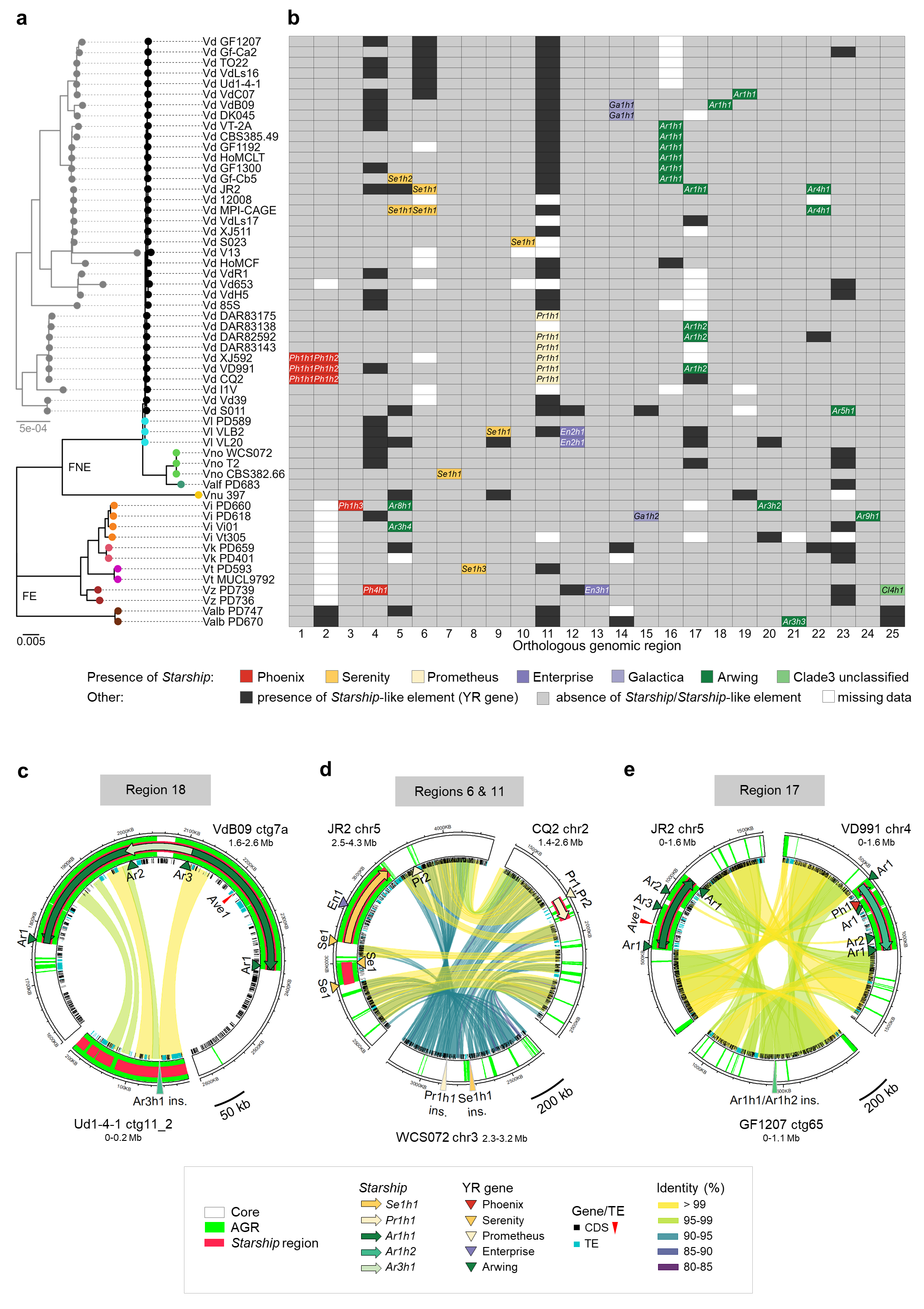

### Supplemental Figure 5

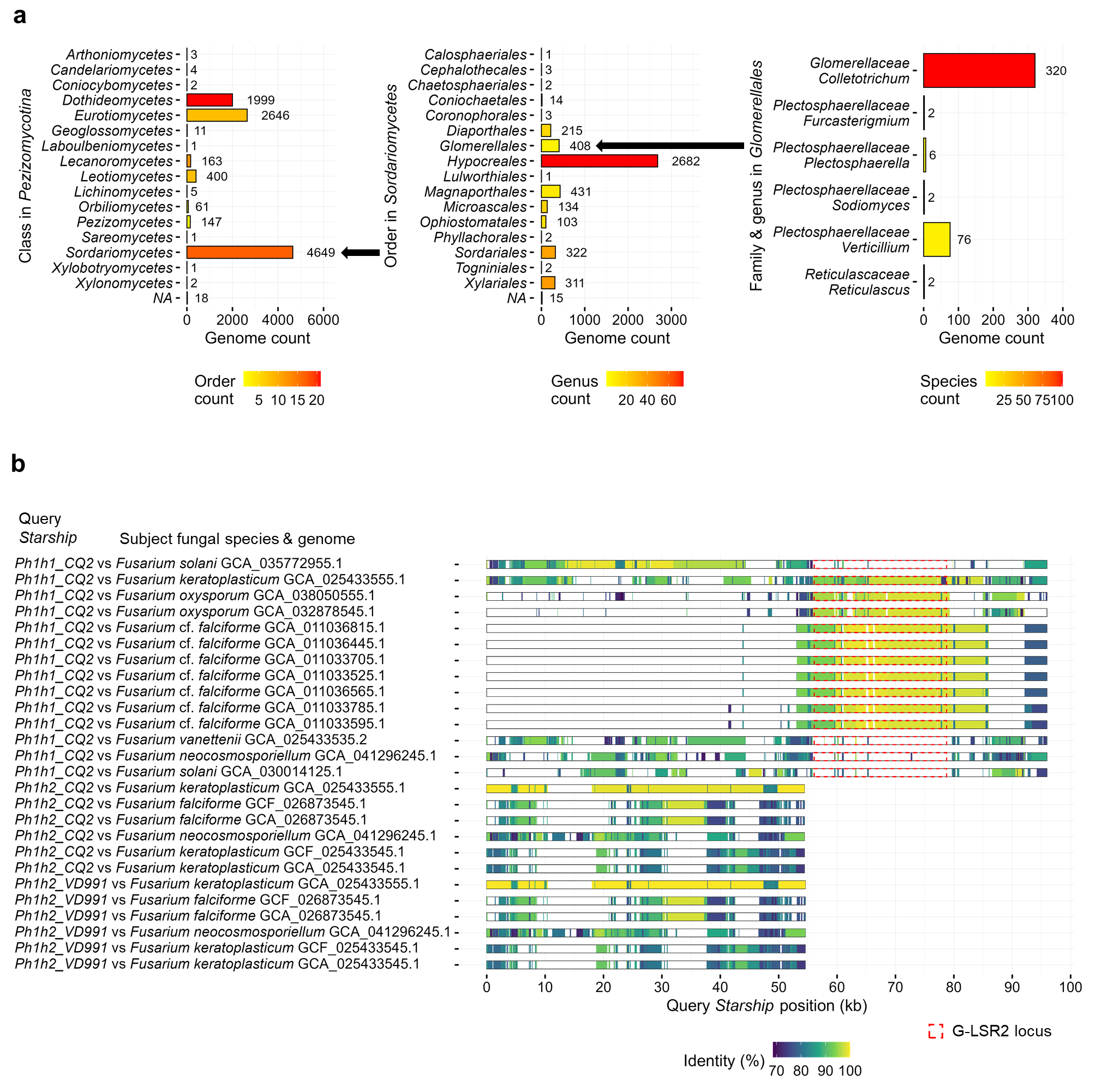

### Supplemental Figure 6

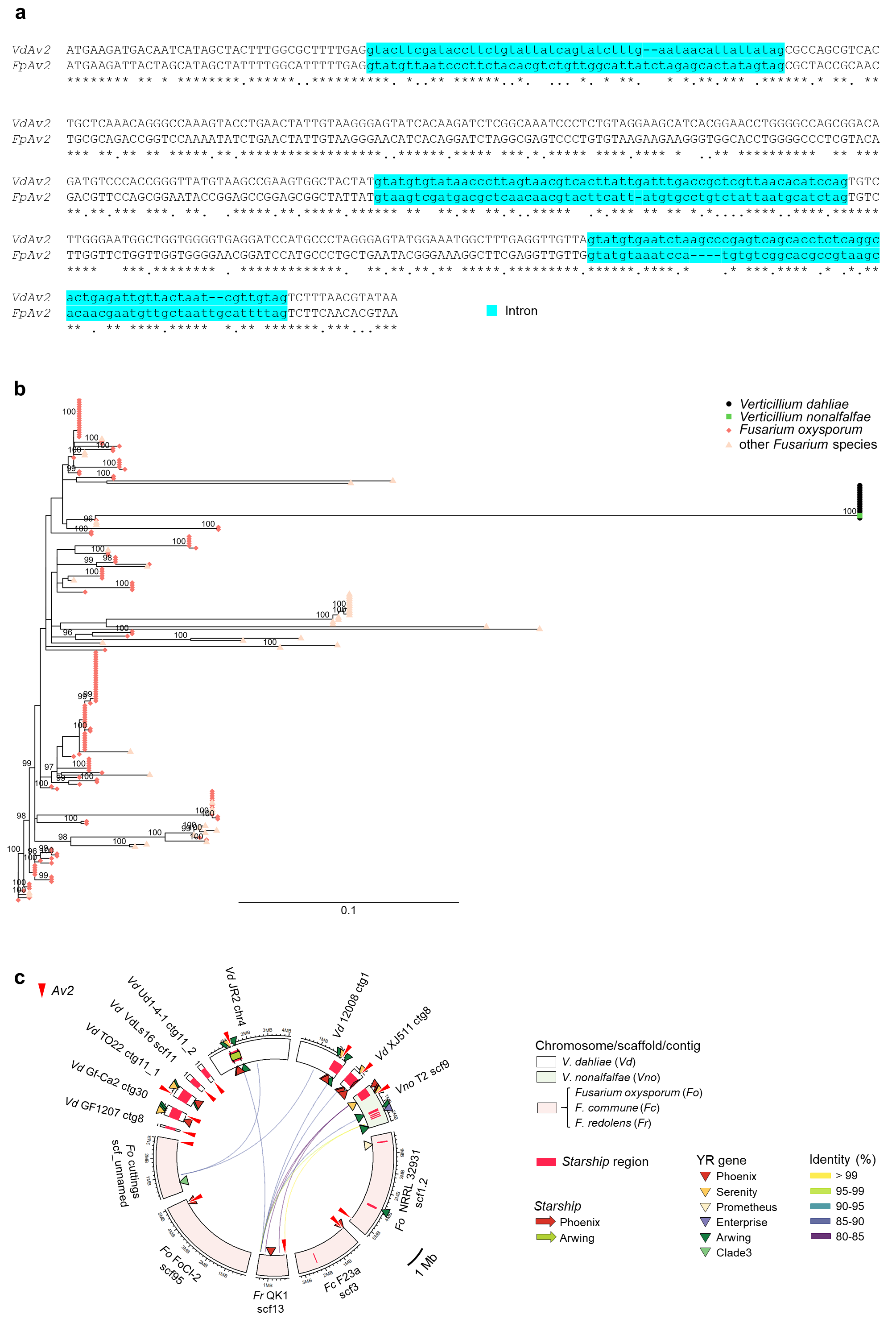

### Supplemental Figure 7

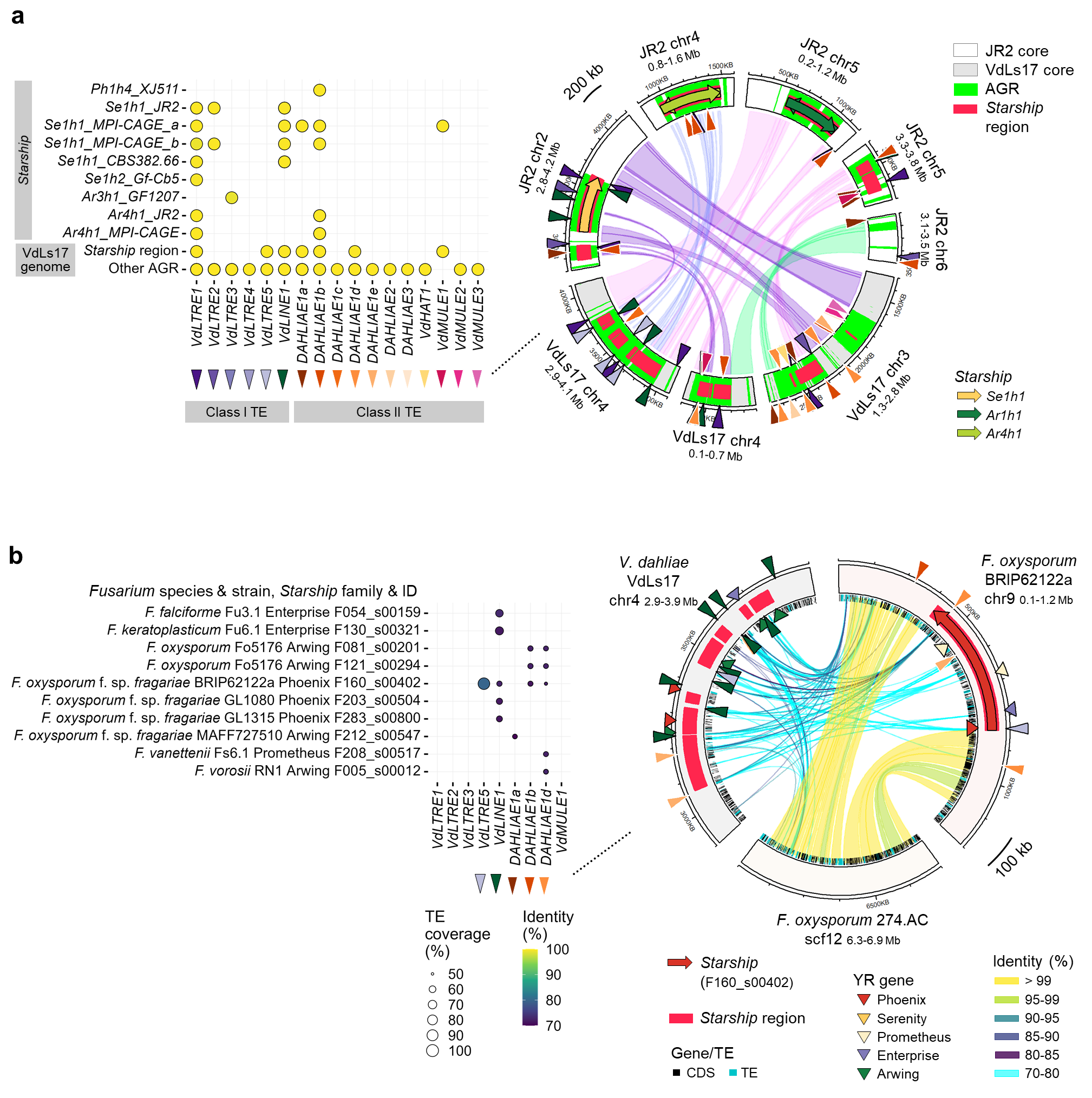

### Supplemental Figure 8

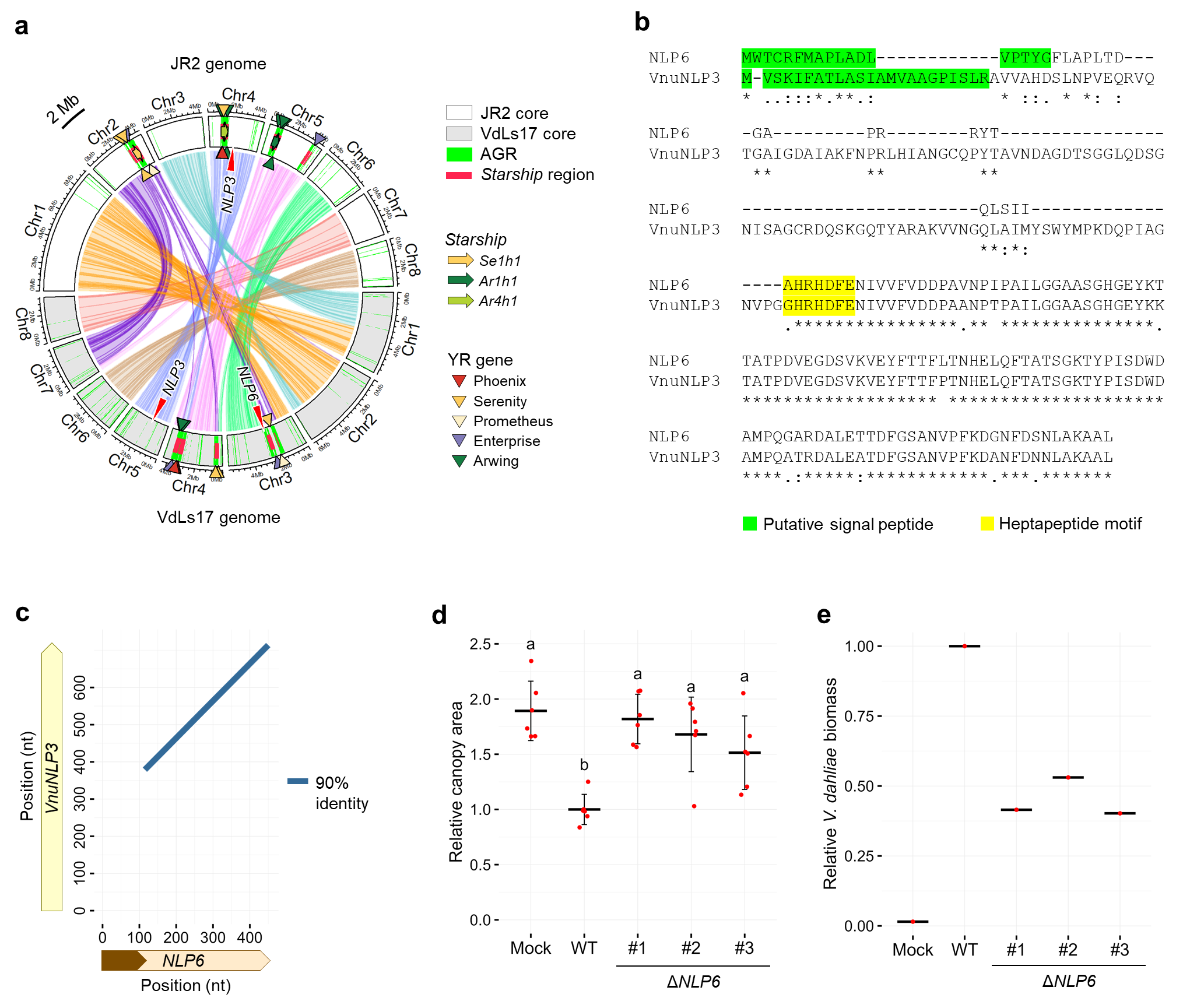
